# A shared ensemble in the prelimbic cortex links impulsivity and anxiety-like behavior

**DOI:** 10.64898/2025.12.25.696439

**Authors:** Rosalie E. Powers, Karla J. Galvan, Daniel E. Calvo, Ricardo Sosa Jurado, Travis M. Moschak

## Abstract

Mental health disorders often share overlapping behavioral and neural features, yet it remains unclear why these relationships emerge or whether they reflect common underlying neural processes. To explore this, we recorded prelimbic (PL) activity using endoscopic calcium imaging as rats completed a battery of tasks assessing impulsivity, distress tolerance, anxiety-like behavior, incentive salience (Pavlovian conditioned approach), and sensation-seeking in a novel environment (locomotor activity). By collecting neural and behavioral measures across all tasks within each animal, we were able to investigate whether PL activity tracked processes that were either unique to individual behaviors, shared across multiple behaviors, or both. We found that PL activity was significantly predictive of each of the behaviors except locomotor activity. Subsequent analyses revealed shared behavioral structure across tasks, with one latent dimension reflecting high impulsivity and low anxiety-like behavior. Individual variability across this particular dimension was strongly predicted by a neural structure comprising shared PL activity across four of the behaviors. Furthermore, this relationship was driven by a subset of PL neurons that shared patterns of activity across multiple tasks, forming a shared neural ensemble. Importantly, animals with a greater tendency to share neural ensembles exhibited a stronger link between high impulsivity and low anxiety-like behavior. These findings suggest that a small, shared ensemble of PL neurons tracked activity across multiple clinically-relevant behaviors to predict an approach/avoidance phenotype characterized by high impulsivity and low anxiety. This points toward a targetable neural population that may help explain why diverse psychiatric symptoms often co-occur within individuals.

## Introduction

Mental health disorders (MHDs) profoundly impact not only the lives of over 1 billion individuals living with these conditions, but also their families, communities, and society at large^1,2^. Although categorized as individual disorders, such conditions are often comorbid with each other^3–5^ and frequently share underlying behavioral phenotypes such as impulsivity^6–9^, anxiety^3,10–12^, sensation-seeking^13–16^, distress tolerance^17–20^ and incentive salience^21–23^. Together, these findings demonstrate that transdiagnostic behaviors are often at the core of many MHDs.

MHDs also frequently share dysfunction in overlapping neural substrates, with notable examples being the default mode network and salience network^24–26^. In addition to the MHDs themselves, many of the underlying behavioral phenotypes mentioned above also exhibit overlapping neural substrates^27–31^, and these neurobehavioral profiles can differentially categorize individuals in a several MHDs^31,32^. Although the shared neural networks implicated in MHDs and their underlying behavioral phenotypes are well established, it is unknown if this overlap is a general feature of these anatomical pathways or if it relies on a distinct subset of neurons that form a shared neural ensemble across behaviors and disorders. Determining the role of such ensembles is important, as preclinical work demonstrates that neural ensembles are critically implicated in models of substance use disorder^33,34^, major depressive disorder^35^, and post-traumatic stress disorder^36,37^. Furthermore, targeting these neural ensembles are likely to play an important role in future therapeutics^38,39^. However, despite the known overlap in behavioral and neural substrates across MHDs, no study has investigated whether such ensembles are shared across distinct clinically-relevant behaviors.

One of the most important nodes in the default mode and salience networks mentioned above is the anterior cingulate cortex, which is homologous to the prelimbic cortex (PL) in rodents^40^. The PL has been implicated in preclinical models of MHDs including substance use disorders^41,42^, major depressive disorder^43,44^, anxiety disorders^45,46^ and eating disorders^47,48^ among others. It also plays an important role in rodent models of behavioral phenotypes such as impulsivity^49,50^, anxiety^51,52^, sensation-seeking^53^, distress tolerance^54^, and incentive salience^55,56^. Several preclinical studies have demonstrated both the presence and absence of shared variance across many of these behaviors^57–59^ and suggested that these interactions (or lack thereof) may drive the multifaceted nature of MHDs^60,61^. However, to our knowledge no studies have determined whether PL ensembles are shared across clinically relevant behavioral phenotypes.

To address this gap, we recorded neural activity from the PL in rats using endoscopic calcium imaging as they completed five distinct behavioral tasks designed to assess impulsivity, anxiety-like behavior, incentive salience (Pavlovian conditioned approach), distress tolerance, and sensation-seeking (locomotor response to a novel environment). By collecting both neural and behavioral data across all five tasks in the same rats, we aimed to investigate whether patterns of activity in the PL tracked processes that were either 1) unique to individual behaviors, 2) reflected shared neural signals across behaviors, or 3) a combination of the two. We were particularly interested in whether individual neurons would show consistent patterns of activation across multiple tasks, suggesting common neural ensembles that underlie different features of psychiatric risk.

## Methods

A detailed description of all procedures and analyses is provided in the Supplement.

### Surgery

Long Evans rats (19 females, 19 males, 8-10 weeks old) were obtained from Envigo (St. Charles, MO). We infused a viral construct encoding GCaMP6s (Addgene) and implanted a GRIN lens (Inscopix) into the PL (**Fig. 2A**). Following 1 week recovery, animals began behavioral training (see below). 6-8 weeks following the initial surgery, we implanted a baseplate over the lens that was optimally aligned to visualize neural activity.

### Behavior

Following surgical recovery, rats underwent a battery of tasks designed to assess impulsivity, distress tolerance, incentive salience (Pavlovian Conditioned Approach), anxiety-like behavior, and locomotor activity. Animals were food-restricted during the impulsivity and distress tolerance tasks and were put on ad libitum during the remaining tasks.

Impulsivity (IMP; **Fig. 1A**). Animals initially trained to press a lever for a sucrose pellet when a cue light was present. The IMP task was composed of a pre-cue period, cue period, and post-cue period (adapted from^49,62,63^). If the lever was pressed during the cue period (when the cue light was illuminated), a sucrose pellet was delivered. If the lever was pressed during either the pre-cue or post-cue periods, no pellet was delivered, and a brief white noise burst was emitted. Early errors were defined as lever presses occurring before cue light illumination (pre-cue period), and impulsivity was quantified as the number of early errors divided by the total number of responses.

**Figure 1.**
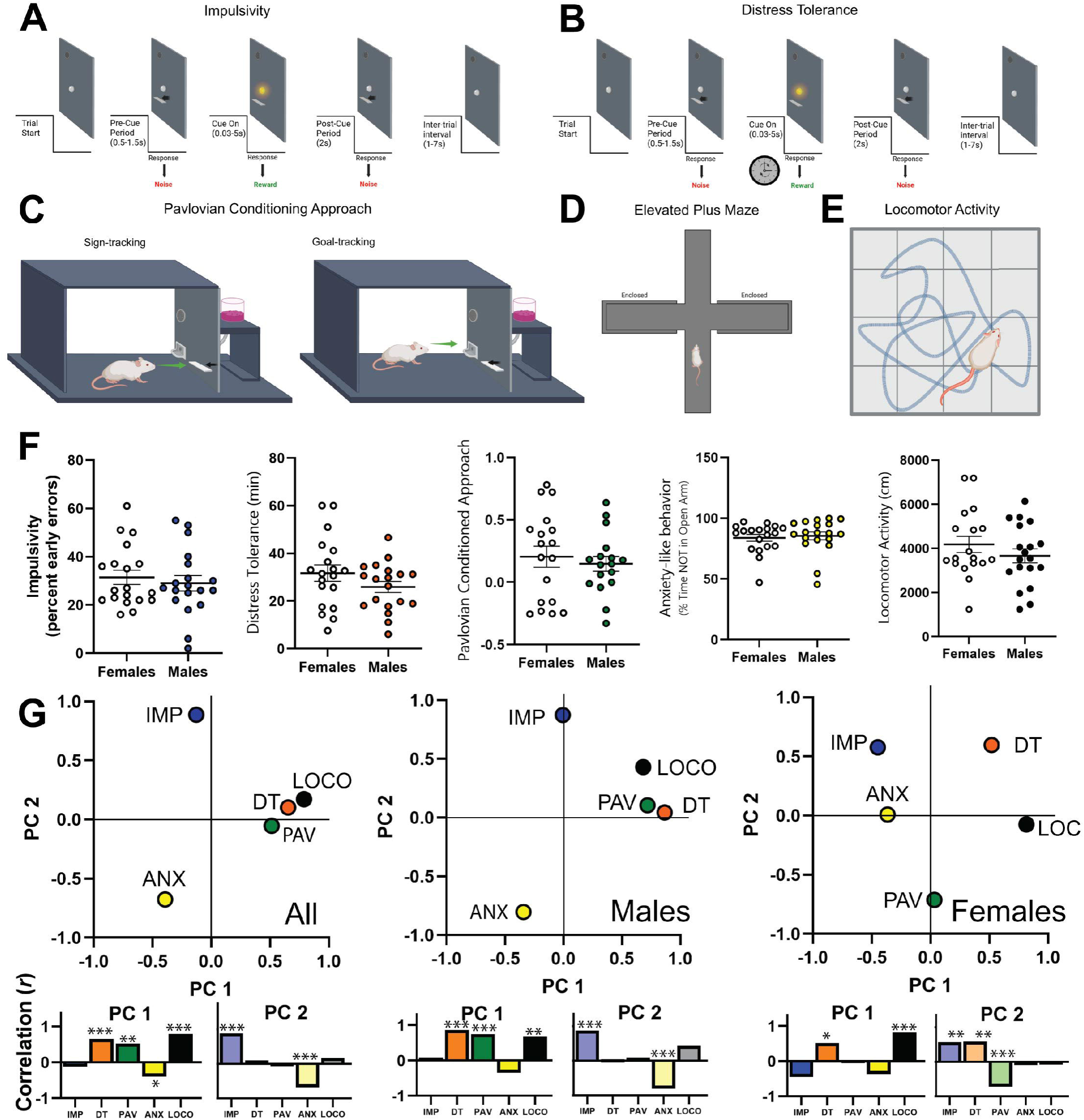
Behavior. **A-E)** Behavior tasks (see methods for details). **F)** Behavioral profiles for each task. There were no differences between males and females (**Table S1**). **G)** Principal component analysis is shown for all animals together as well as separately for males and females. *Left.* Overall analysis. The first principal component had strong loadings from locomotor activity (0.789) and distress tolerance (0.654). The second principal component had strong loadings from impulsivity (0.887) and anxiety-like behavior (−0.676). PCA for males (*middle*) closely tracked the overall PCA, while females (*right*) were similar but deviated in anxiety-like behavior. Nonetheless, there were no differences in male and female scores when they were independently loaded onto the overall PCA (PC1: t(36) = 1.53, p = 0.134; PC2: t(36) = 0.53, p = 0.602). Variance explained: Overall PC1: 34.58%, PC2: 20.86%; Male PC1: 48.39%, PC2: 20.82%; Female PC1: 25.72%, PC2: 23.74%. The areas under each plot depict correlations of the PCs with the raw behavioral data (excluding estimated data, see **Table S2**) * p < 0.05, ** p < 0.01, *** p < 0.001

Distress Tolerance (DT; **Fig. 1B**). The DT task (adapted from^64^) was similar to the IMP task with a pre-cue period, cue period, and post-cue period. Unlike the IMP task, during the DT task the cue light duration progressively decreased following each response until it was impossible for the rat to obtain a correct response. DT was defined as the duration of time that elapsed before the rat ceased to respond on the lever (we specifically defined DT as the timepoint when the rat obtained 8 response omissions out of the past 10 trials).

Pavlovian Conditioned Approach (PAV; **Fig. 1C**). The Pavlovian conditioned approach task was used to assess incentive salience (i.e. the motivational value given to cues associated with rewards). Rats underwent a 5-day PAV training protocol prior to recording. On each trial, a retractable lever serving as a cue was extended for 8 seconds, followed immediately by pellet delivery. Lever interactions and food cup entries were recorded. A Pavlovian conditioned approach index was calculated to quantify preference for the lever (sign-tracking) or preference for the food cup (goal-tracking) using approach probability, response bias, and initial preference (modified from^65^).

Anxiety-like behavior (ANX; **Fig. 1D**). The elevated plus maze (EPM) was used to assess anxiety-like behavior. Animals were placed in the direct center of the maze, facing an open arm and allowed to explore for 10 minutes. Behavior was recorded via the number of infrared beam breaks at the junction of each arm. Percent time not in the open arms was calculated as a measure of anxiety-like behavior.

Locomotor Activity (LOCO; **Fig. 1E**). Locomotor activity was measured in an open-field arena under dim light. Animals were allowed to explore the arena freely for 10 minutes. Infrared beam breaks were used to record total distance travelled.

### Calcium Imaging

Calcium imaging was recorded during tasks using the UCLA V4 Miniscope (Open Ephys, Lisbon, Portugal). During each task, the camera was attached to the implanted baseplate on the animal’s skull and removed immediately after the task was over. Recordings were captured for each task, with 5-minute breaks every 10 minutes to reduce photobleaching. Raw video files were downsampled and processed using CaImAn^66,67^ (**Fig. 2B**). Putative neurons within the videos were identified using an algorithm that highlights candidate neurons based on the spatiotemporal signature of each calcium signal. A separate algorithm coregistered neurons across separate tasks.

**Figure 2.**
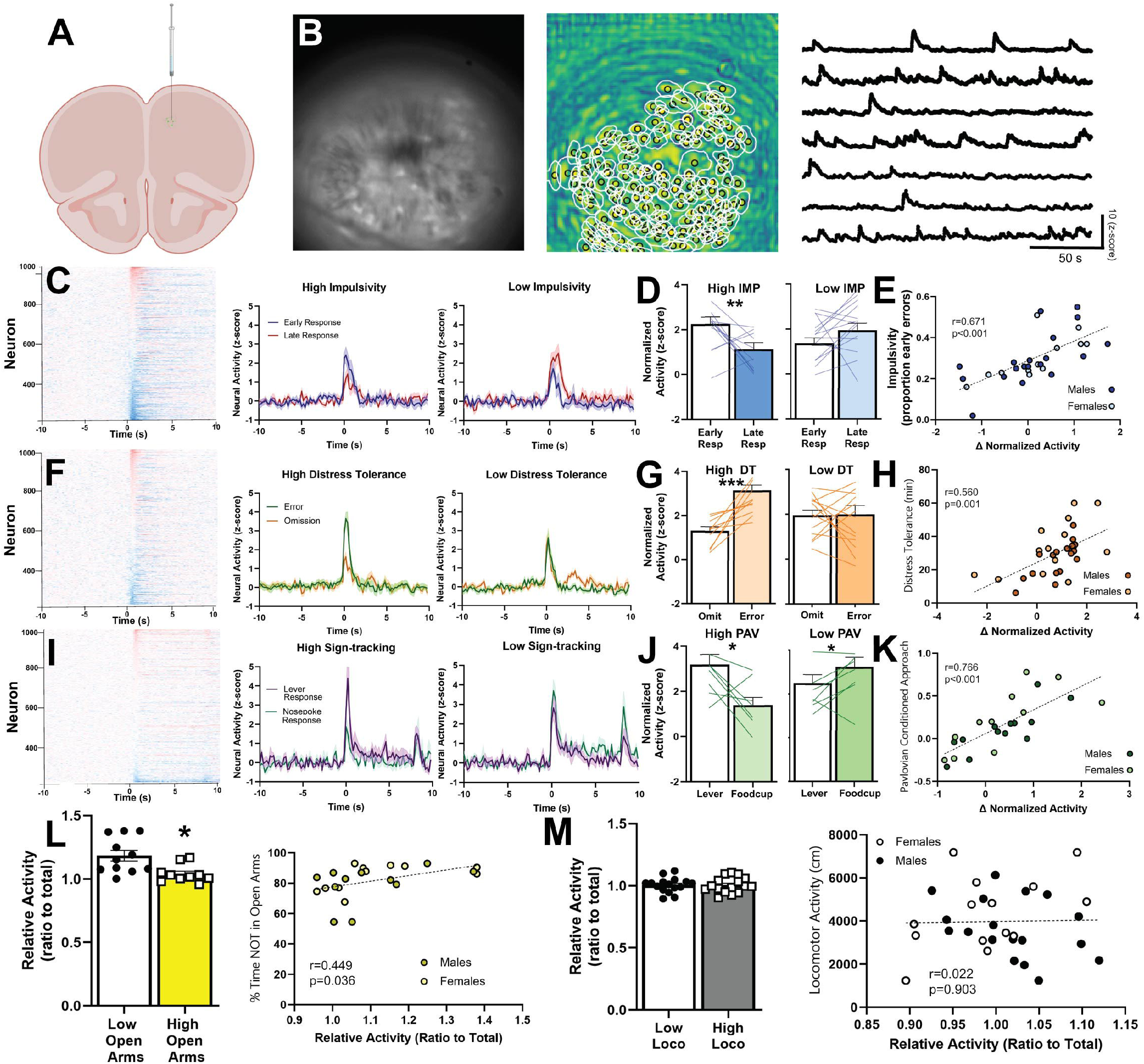
Prelimbic activity during behavior. **A)** GCaMP6s was infused into the prelimbic cortex, followed by a GRIN lens. **B)** *Left.* Frame of a sample video using the miniscope. *Middle.*Components classified as putative cells by CaImAn. *Right.* Calcium traces of putative neurons captured by CaImAn. **C)** PL activity during the impulsivity task. *Left.* Perievent activity to trial initiation for each neuron. *Middle and right.* PL activity in neurons that were excited by trial initiation. Animals with high impulsivity had a more pronounced increase in PL activity when they subsequently chose an early response, while animals with low impulsivity showed the opposite pattern. In this and other figures animals were divided into low and high groups via median split. **D)** High impulsive rats had more activity to the lever when they subsequently chose an early response (Low IMP: t(13) = 1.57, p = 0.141; High IMP: t(12) = 3.06, p = 0.0099). **E)** The total pattern, including inhibited neurons, as a correlation. Animals with a greater change in activity (early response – late response) had higher impulsivity. **F)** PL activity during the distress tolerance task. *Left.* Perievent activity to trial initiation for each neuron. *Middle and right.* PL activity in neurons that were excited by trial initiation. Animals with high distress tolerance had a more pronounced increase in PL activity when they subsequently made a response, while animals with low distress tolerance did not. **G)** High DT rats had more activity when they subsequently made a respondse (High DT: t(13) = 5.74, p < 0.001; Low DT: t(13) = 0.09, p = 0.9296). **H)** The total pattern, including inhibited neurons, as a correlation. Animals with a greater change in activity (error response – omission) had higher distress tolerance. **I)** PL activity during the Pavlovian conditioned approach task. *Left.* Perievent activity to trial initiation for each neuron. *Middle and right.* PL activity in neurons that were excited by trial initiation. Animals with high sign-tracking had a more pronounced increase in PL activity when they subsequently approached the lever, while animals with low sign-tracking showed the opposite pattern. **J)** High sign-trackers had more activity when they subsequently interacted with the lever; low sign-tracker had more activity when they subsequently interacted with the food cup (High PAV: t(8) = 2.91, p = 0.0197; Low PAV: t(8) = 2.41, p = 0.042). **K)** The total pattern, including inhibited neurons, as a correlation in. Animals with a greater change in activity (lever response – nosepoke response) had higher sign-tracking. **L)** In the elevated plus maze, animals that spent more time in the open arms exhibited less PL activity in the open arms (t(20) = 3.20, p = 0.047). **M)** There were no relationships between PL activity and locomotor activity (t(30) = 0.02, p = 0.980). * p < 0.05, ** p < 0.01, *** p < 0.001.

### Data analysis

Neural activity was classified according to our previous methods^49,51,68^. A single neural data point per animal per task was derived from either averaged differential perievent neural activity in trial-based tasks (aligned to trial start in the IMP, DT, and PAV tasks) or activity-based neural activity (e.g. when animal was in open arm, or when animal was moving) in the ANX and LOCO tasks. Neural and behavioral data points were then correlated for each behavior to establish the individual relationship between neural activity and behavior. To investigate shared variance between the different behaviors, principal component analysis (PCA) was run on data from all five behaviors to derive underlying latent variables for both behavior and neural activity. Then, to determine if a common neural-behavior relationship spanned multiple behaviors, we ran a canonical correlation analysis (CCA) as per^69^. Finally, we investigated whether shared neural ensembles were the drivers of shared predictive relationships between neural activity and behavior. To do so, we first assessed whether there were any “dominant neural patterns” between each set of two behaviors. An example of a dominant pattern would be if a statistically significant majority of neurons that were excited in one task were then inhibited in another task. Individual neurons that followed the dominant neural pattern between two tasks were considered to be part of a shared neural ensemble for those tasks. Once all neurons were classified as either part of a shared neural ensemble (‘shared’) or not (‘unshared’), we separately ran PCA and CCA on ‘shared’ and ‘unshared’ neural populations. Finally, we used a moderation analysis^70^ to investigate whether individual differences in the proportion of neurons that were shared (‘neural overlap’) moderated the predictive relationships between behaviors across tasks.

## Results

### Distinct latent variables underlie shared variance in behavior

Following initial training, we assessed each of the behaviors described in **Fig. 1A-E**. We found no sex differences in behavior (**Fig. 1F, Table S1**) and used PCA to determine if there were latent variables underlying the different behavioral phenotypes. We found that the behavior data separated into two distinct principal components with strong loadings from LOCO and DT (PC1) or IMP and ANX PC2, see(**Fig. 1G *left*, Table S2**). The first principal component had strong loadings from LOCO and DT. The second had strong loadings from IMP and ANX. We saw no differences in male and female scores when they were independently loaded onto the overall PCA, which suggests the overall PCA is valid for capturing both sexes (**Fig. 1G *middle, right*, Table S2**). In total, these data suggest that behavioral variation in LOCO and DT forms a distinct latent variable from IMP and ANX.

### Prelimbic activity tracks multiple behaviors individually

Next, we determined if PL activity predicted behavior in each of the behaviors individually (i.e. not in combination with other behaviors). We found that PL activity significantly predicted behavioral patterns in four of the five behaviors we assessed. For impulsive animals in the IMP task, we found that both excited and inhibited neurons exhibited a stronger response to trial start when the animal subsequently made an impulsive response, and that this pattern significantly predicted IMP (**Fig. 2C-E, S1A-B**). Similarly, animals with high distress tolerance in the DT task had stronger excited and inhibited activity to trial start when they subsequently made a response (**Fig. 2F-H, S1C-D**) and animals exhibiting high Pavlovian conditioned approach (PAV, often referred to as high sign-tracking) in the Pavlovian task had stronger activity to trial start before making sign-tracking responses (**Fig. 2I-K, S1E-F**). Furthermore, animals with higher ANX had significantly higher PL activity when in the open arms (**Fig. 2L**). Finally, PL activity did not predict locomotor activity (**Fig. 2M**). These findings demonstrate that PL activity is a significant predictor in multiple distinct behaviors that are directly relevant to MHDs. We next sought to investigate if common elements of this neural activity served to predict shared variance across the behaviors we tested.

### Latent variables in prelimbic activity collectively predict impulsivity and anxiety-like behavior

Using principal component analysis, we found that PL data for each behavior clustered into two distinct principal components (**Fig. 3A *left*, Table S3**). The first principal component had strong loadings from PL activity during DT and IMP, while the second had strong loadings from PL activity during ANX, PAV, and LOCO. We saw no differences in male and female scores loaded onto the overall PCA, which suggests the overall PCA is valid for capturing both sexes (**Fig. 3A *middle*, *right*, Table S3**). Together, these data suggest that PL activity during DT and IMP forms a distinct cluster from that seen during ANX, PAV, and LOCO.

**Figure 3.**
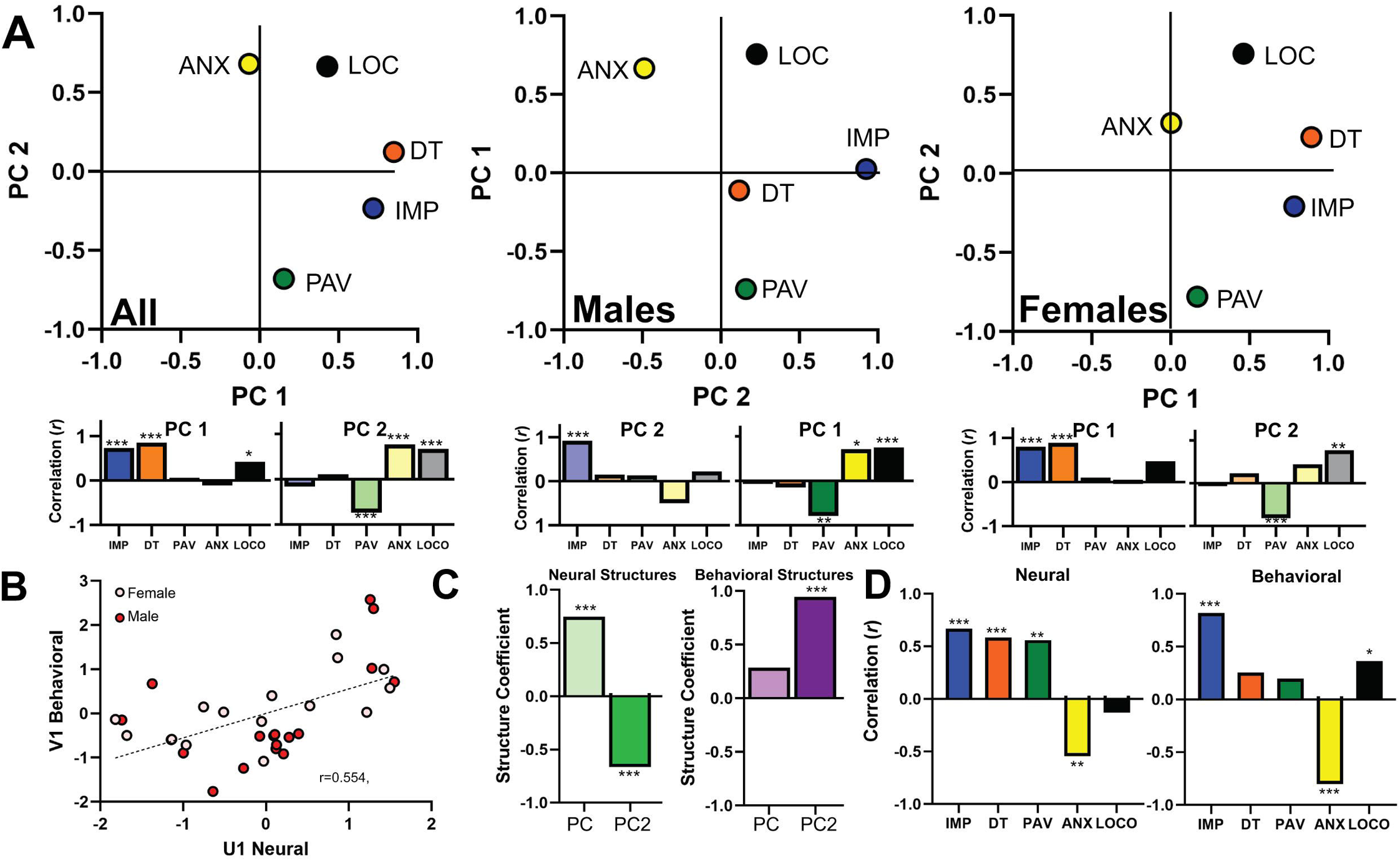
Prelimbic activity across multiple behaviors. **A)** Principal component analysis is shown for all animals together as well as separately for males and females. On PC1, strong loadings were seen for PL activity during distress tolerance (0.850) and impulsivity (0.720). On PC2, moderately strong loadings were seen for PL activity during elevated plus maze (0.682), Pavlovian conditioned approach (−0.679), and locomotor activity (0.665). PCA for females closely tracked the overall PCA, while males were similar but deviated in distress tolerance. Nonetheless, we saw no differences in male and female scores loaded onto the overall PCA (PC1: t(31) = 0.42, p = 0.676; PC2: t(31) = 0.22, p = 0.825; note that in the central graph (males) the PC axes are switched, see Supplement for rationale). Variance explained: Overall PC1: 30.39%, PC2: 27.43%; Male PC1: 34.18%, PC2: 23.33%; Female PC1: 36.29%, PC2: 24.44%. The areas under each plot depict correlations of the PCs with the raw behavioral data (excluding estimated data, see **Table S2**). **B)** To determine the collective relationship between neural activity and behavior across all behaviors, we used a Canonical correlation analysis. We found that the first pair of canonical variates were significantly correlated. **C)** The neural covariate had significant positive influence from PC1 and negative influence from PC2 (PC1: r = 0.748, p < 0.001; PC2: r = −0.663, p < 0.001, *left*), while the behavioral covariate had significant positive influence from PC2 (PC1: r = 0.284, p = 0.109; PC2: r = 0.944, p < 0.001, *right*). **D)** Finally, we correlated the neural and behavioral covariates with the raw neural and behavioral scores (excluding estimated data). We found significant positive contributions from neural activity during the IMP, DT, and PAV tasks and a significant negative contribution from neural activity during the EPM task (IMP: r = 0.669, p < 0.001; DT: r = 0.584; p < 0.001 ; PAV: r = 0.559, p = 0.016; ANX: r = −0.545, p = 0.035; LOCO: r = −0.128; p = 0.499, *left*). This combined activity predicted the behavioral covariate, which had significant positive contributions from the IMP, EPM and LOCO tasks (IMP: r = 0.819, p < 0.001; DT: r = 0.253, p = 0.155; PAV: r = 0.198, p = 0.297; ANX: r = −0.803, p < 0.001; LOCO: r = 0.362, p = 0.042, *right*). * p < 0.05.

Having identified latent variables in both the neural and behavioral data, we next used canonical correlation analysis (CCA) to investigate whether the shared variance across the two neural principal components in **Fig. 3A** collectively predicted shared variance across the two behavioral principal components in **Fig. 1G**. We found that shared neural variance significantly predicted a portion of the shared behavioral variance in the data (**Fig. 3B, S2A**). To further understand the specific components of this relationship, we investigated how the neural structures (neural PC1 and PC2) related to the shared neural activity (canonical variate U_1_) and how the behavioral structures (behavioral PC1 and PC2) related to the shared behavioral activity (canonical variate V_1_). The two neural PCs both played a role in the shared variance, as each had an opposing impact on the shared neural activity (**Fig. 3C *left***). Conversely, only the second behavioral PC was related to the shared behavioral activity (**Fig. 3C *right***). Thus, the data suggest that the two neural PCs independently have opposing predictive influence on behavioral PC2, while behavioral PC1 is not driven by shared variance in the neural activity. To understand how this related to our specific behavioral measures, we correlated each canonical variate with their respective raw neural or behavioral indices (excluding any estimated data used in the PCA analysis, see Supplement for details). Our neural canonical variate (U_1_) positively correlated with neural activity in the IMP, DT, and PAV tasks while correlating negatively with neural activity in the ANX task (**Fig. 3D *left***). Our behavioral canonical variate (V_1_) had a strong positive association with IMP and a strong negative association with ANX, as well as a weaker positive association with LOCO (**Fig. 3D *right***). In total, shared PL activity across all behaviors except for locomotor activity collectively predicted high IMP and low ANX, which together formed the second principal component in our behavioral data. Conversely, our first behavioral principal component, which was strongest for DT, PAV, and LOCO, was not predicted by shared neural activity. Thus, in the PL these latter behaviors appear to be predicted solely by the individual neural activity specific to the relevant behavior (see **Fig. 2**).

### A subset of prelimbic neurons formed a shared neural ensemble that drove the common relationships between neural activity and behavior

To investigate whether the shared predictive relationship described in **Fig. 3B** was driven by those neurons that shared similar patterns of activity across behavioral tasks (i.e. a ‘shared neural ensemble’), we first assessed whether there were any dominant patterns of activity shared across each pair of tasks. For this, we determined phasic activity (i.e. if a neuron was significantly excited or inhibited during an event) for each rat across each pair of behaviors. We first investigated the tasks that had a trial-based structure (IMP, DT, and PAV). When looking at these trial-based tasks, we found that phasic PL neurons during the IMP and DT task were significantly more likely to share a ‘positive’ pattern of activity (e.g. “excited” neurons in IMP tended to also be “excited” during DT and “inhibited” neurons in IMP tended to also be “inhibited” during DT; **Fig. 4A**). A similar but weaker effect was seen for DT and PAV neurons (**Fig. 4A**). Next, we investigated the relationship between ANX and the trial-based tasks. We found that phasic PL neurons during the IMP and ANX tasks showed a ‘negative’ pattern of activity (e.g. “excited” neurons in IMP tended to be “inhibited” during ANX and vice versa; **Fig. 4B**). In total, there were three pairs of tasks exhibiting a statistically significant dominant pattern: IMP and DT (positive, **Fig. 4A**), DT and PAV (positive, **Fig. 4A**), and IMP and ANX (negative, **Fig. 4B**). As a result, we classified “shared” neurons as those phasic neurons that followed the dominant pattern within each set of tasks, while “unshared” neurons were those phasic neurons that did not. Averaging across all datasets, shared neurons represented 15.53% of all neurons, while unshared neurons represented 32.59%.

**Figure 4.**
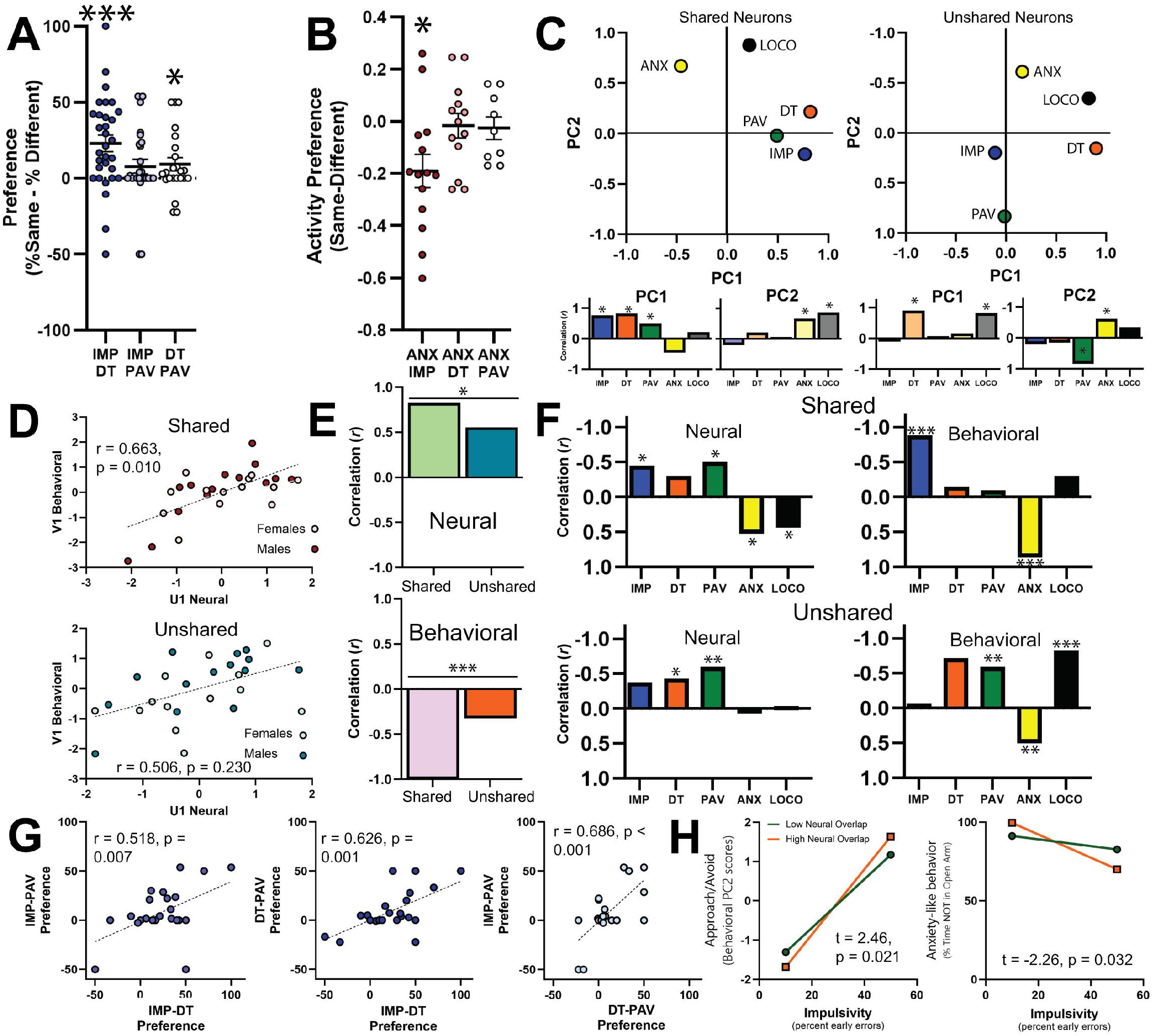
Neurons that share patterns of activity across multiple tasks drive the relationship to behavior. **A)** In trial-based tasks, phasic neurons that were coregistered across the IMP and DT tasks as well as the DT and PAV tasks had a significant preference for the same pattern of neural activity (IMP/DT: t(30) = 4.257, p < 0.001; DT/PAV: t(22) = 2.155, p = 0.042). **B)** Phasic neurons coregistered across the EPM task and IMP task significantly preferred to have the opposite pattern of activity (IMP/ANX: t(13) = −2.967, p = 0.011). **C)** Principal component analysis for shared (*left*) and unshared (*right*) neurons. On shared neurons PC1, strong loadings were seen for PL activity during DT (0.822) and IMP (0.768). On PC2, strong loadings were seen for PL activity during LOCO (0.877) with moderately strong loadings from EPM (0.671). On unshared neurons PC1, strong loadings were seen for PL activity during DT (0.901) and LOCO (0.828). On PC2, a strong loading was seen for PAV (0.832) and a moderately strong loading from EPM (−0.611). Shared neurons variance explained: PC1: 35.78%, PC2: 25.70%; Unshared neurons variance explained: PC1: 34.16%, PC2: 23.18%. The areas under each plot depict correlations of the PCs with the raw behavioral data (excluding estimated data, **Table S4**). **D)** Canonical correlation analysis for shared (*top*) and unshared (*bottom*) neurons. The first pair of canonical variates were significantly correlated in shared neurons, but not unshared neurons. **E)** Furthermore, both neural (*top*) and behavioral (*bottom*) covariates had a significantly stronger correlation with the overall analysis in shared rather than unshared neurons (neural comparison between r_shared_ and r_unshared_, Fisher’s Z: 1.867, p = 0.031; behavioral comparison between r_shared_ and r_unshared_, Fisher’s Z: 11.03, p < 0.001). **F)** Finally, we correlated the neural and behavioral covariates with the raw neural and behavioral scores (excluding estimated data). We found significant negative contributions from neural activity in shared neurons during the IMP, and PAV tasks and a significant positive contribution from neural activity during the EPM and LOCO tasks (*top left*; **Table S5**). This combined activity predicted the behavioral covariate, which had significant negative contributions from the IMP and EPM tasks (*top right*; **Table S5**). These patterns were markedly similar to the overall analysis. Conversely, the unshared neurons differed, most notably in EPM neural activity and DT and PAV behavior (*bottom*; **Table S5**); note that in each set of graphs the Y axes are inverted, see Supplement for rationale **G)** Individual differences in preference patterns of activity in one set of tasks predicted the same preference across the other sets of tasks. **H)** Neural overlap moderated the relationship between impulsivity and behavioral PC2 (*left*; interaction term: t(27) = 2.45, p = 0.021) and anxiety-like behavior (*right*; interaction term: t(27) = 2.26, p = 0.032). * p < 0.05, ** p < 0.01, *** p < 0.001

Next, we examined PCA and CCA separately in shared and unshared neural populations. Following PCA, we found that each subset of neurons had a similar structure to that of the overall dataset (with a notable exception of PAV for the shared neurons and IMP for the unshared neurons; compare shared [**Fig. 4C *left***] and unshared [**Fig. 4C *right***] with overall dataset [**Fig. 3A *left***]). However, when we conducted the CCA separately for each population of neurons, we found that the first pair of canonical variates were significantly related for shared neurons but not unshared neurons (**Fig. 4D**). To directly compare the shared and unshared neurons as they related to the overall analysis, we first took the neural variates for each set (U_1-shared_ and U_1-unshared_) and correlated them with the neural variate for the overall analysis (U_1-overall_). Then we compared these correlations to each other using Fisher’s r-to-z test. We did the same for the behavioral variates. We found that the shared neurons had a significantly stronger relationship to both neural and behavioral canonical variates of the overall CCA than the unshared neurons (**Fig. 4E, S3**). For the shared neural population, raw neural activity and behavior exhibited strikingly similar relationships to the canonical variates as that seen in the overall CCA (compare **Fig. 4F *top*** to **Fig. 3D**). This was not seen for unshared neurons (**Fig. 4F *bottom***). In total, the similarities seen between the shared CCA analysis and the overall analysis (and lack thereof in the unshared neurons) suggests that the subset of neurons that share patterns of neural activity across tasks are the ones that drive the shared predictive relationships between neural activity and behavior.

### Individual differences in neural overlap moderate the relationship between impulsivity and anxiety

Notably, there were wide individual differences in the degree that rats shared patterns of neural activity across tasks (**Fig. 4A, B**). Interestingly, for all the trial-based tasks we found that individual differences in shared patterns of activity for one set of tasks predicted individual differences in shared patterns of activity in other sets of tasks (**Fig. 4H**), suggesting that some rats have a general propensity to share patterns of PL activity across tasks while others don’t. To explore if this tendency towards shared neural ensembles would impact behavior, we calculated an overall “neural overlap” index for each rat by averaging “neural preference” across the lever-based tasks and the ANX task (see Methods and Supplement for details). Then, we investigated the influence of “neural overlap” on the relationship between neural activity and behavior, particularly for our relevant component behaviors of PC2, i.e. IMP and ANX. Using a moderation analysis with neural overlap as the moderator, we found that IMP was significantly more predictive of behavioral PC2 in animals with high neural overlap than in animals with low neural overlap (**Fig. 4I *left***). Similarly, we further found that IMP was significantly more negatively predictive of ANX itself in animals with high neural overlap than in animals with low neural overlap (**Fig. 4I *right***). Together, these data demonstrate that animals with a high propensity to share neural ensembles across tasks also had a stronger link between IMP, ANX, and behavioral PC2.

## Discussion

MHDs are often comorbid and share common underlying behaviors and neural activity^3–23^. However, the role that neural ensembles play in the relationships between these behavioral phenotypes is unknown. Here, we investigated behavioral and neural commonalities across five clinically-relevant behaviors. We found shared dimensions that linked many of our tasks both neurally and behaviorally. In particular, we found that shared PL activity across four of the five behaviors (all but locomotor activity) significantly predicted a latent behavioral pattern underlying high impulsivity and low anxiety-like behavior. This relationship was driven by a subset of approximately 16% of PL neurons that shared similar patterns of activity across multiple behaviors (thus forming a shared neural ensemble). Notably, individuals with a greater tendency to share neural ensembles also exhibited a stronger relationship between high impulsivity and low anxiety-like behavior. Together these findings suggest that shared neural ensembles in the PL drive the predictive relationships between these behaviors.

Behaviorally, we identified two latent variables – one associated with locomotor activity and distress tolerance, and the other with impulsivity and low anxiety-like behavior. The first latent variable may reflect motivational persistence in the absence of reward, since animals high in this trait would be expected to have both high distress tolerance and high locomotor activity. Clinical research supports a relationship, as high sensation-seeking individuals are known to persist in risky or effortful contexts despite low payoff and show reduced responsiveness to negative feedback^71–73^. Such individuals may be guided more by internal stimulation thresholds or novelty-seeking than by conventional reward-punishment. Conversely, the second latent variable may reflect an approach/avoid phenotype. The conceptualization of impulsivity and anxiety as opposite poles of an approach/avoid phenotype was originally postulated in Gray’s behavioral activation/inhibition system^74^. Subsequent research both supports^75,76^ and complicates^77–79^ this, as impulsivity and anxiety appear to be just two of many aspects associated with ‘behavioral activation’ and ‘behavioral inhibition’^78,79^. Our current study, which found no raw correlation between impulsivity and anxiety-like behavior, fits with this complicated picture in suggesting that impulsivity and anxiety may share a core ‘approach/avoid’ phenotype, but also retain their own unique aspects.

Neurally, we also identified two latent variables – one associated with neural activity during impulsivity and distress tolerance, and one with neural activity during locomotor activity, anxiety-like behavior, and low sign-tracking. The first latent variable may reflect a neural preference for action: the preference for immediate impulsive response and the preference for persistence in responding. Conversely, the second latent variable appears unrelated to action or inaction per se. Rather, it may be related to behavioral preference when there are multiple options; here, preference for the food cup over the lever and the closed arm over the open arm. Prior clinical work has suggested that the anterior cingulate tracks domain-general comparison between choice^80^, but further work would be needed to verify this in our paradigm.

When we analyzed neural and behavioral activity together, we found that the two neural principal components collectively predicted the latent variable we are calling ‘approach/avoid’ that is associated with impulsivity and low anxiety-like behavior. Notably, a role for the PL (or homologous ACC) in canonical measures of approach/avoid behavior has been previously demonstrated both preclinically^81,82^ and clinically^83,84^. However, our study offers new insight into this relationship by demonstrating that there are two distinct patterns of PL activity feeding into this behavior, and that these neural patterns span multiple individual behavioral measurements including those unrelated to ‘approach/avoid’ behavior. Furthermore, our work demonstrates that the relationship between PL activity and ‘approach/avoid’ behavior is driven by a small ensemble (∼16% of recorded neurons) that shares patterns of activity across multiple behaviors. Several prior studies have demonstrated a role for shared neural ensembles in the PL, especially for learning^85–88^ and reward valuation^89–92^. Our study expands upon this work by demonstrating that PL ensembles are shared across multiple unrelated and clinically relevant behaviors. Furthermore, we also found individual differences in the degree to which rats formed these shared neural ensembles. A recent study found similar individual differences when examining food and cocaine reward^91^, although the study found no relationships between the degree of neural overlap and behavior. Here we found that the degree of neural overlap significantly moderated the relationship that impulsivity had with both our ‘approach/avoid’ latent variable and with anxiety-like behavior itself. In animals with high neural overlap, impulsivity strongly predicted both ‘approach/avoid’ and anxiety-like behavior. In animals with low neural overlap, the relationship was weaker or nonexistent. Put simply, rats with high neural overlap had high behavioral overlap. Although our results are specific to impulsivity and anxiety-like behavior, they suggest that any two seemingly unrelated behaviors may share a stronger relationship than data initially suggest, especially when focusing on those individuals with high neural overlap in a given brain region. Application of this knowledge to other studies may further our understanding of the transdiagnostic role a given phenotype plays across disorders.

In conclusion, our results demonstrate that a shared neural ensemble in the PL tracks activity across multiple unrelated behaviors to predict an ‘approach/avoid’ phenotype comprising high impulsivity and low anxiety. These results suggest that shared neural ensembles provide a critical link between clinically-relevant behavioral phenotypes, and further points toward a targetable neural population that may help explain why diverse psychiatric symptoms often co-occur within individuals.

## Acknowledgements

This work was supported by and National Institute of Health (NIH) grant U54MD007592-28, National Institute on Drug Abuse (NIDA) grant DA045764, and National Institute on Drug Abuse (NIDA) grant DA058653 awarded to T.M.M. The authors would also like to thank Alyssa Lopez, Sofia Grijalva Torres, Ymalay Vega and Marina Smoak for technical assistance as well as Judy Prasad for comments on an earlier draft of the manuscript. Imaging was performed at the Imaging & Behavioral Neuroscience (IBN) Facility Imaging Core at The University of Texas at El Paso, supported by the College of Science and the Office of Research & Innovation. The IBN Facility was supported by the Office of the Director, National Institutes of Health (NIH), under award number C06OD032074. We thank Sivasai Balivada for his assistance. The content is solely the responsibility of the authors and does not necessarily represent the official views of the NIH.

## References

1. Rehm, J. & Shield, K. D. Global Burden of Disease and the Impact of Mental and Addictive Disorders. Curr. Psychiatry Rep. 21, 10 (2019).

2. World Health Organization. Mental health atlas 2024. https://www.who.int/publications/i/item/9789240114487.

3. Vorspan, F., Mehtelli, W., Dupuy, G., Bloch, V. & Lépine, J.-P. Anxiety and Substance Use Disorders: Co-occurrence and Clinical Issues. Curr. Psychiatry Rep. 17, 4 (2015).

4. Hartman, C. A., et al. Anxiety, mood, and substance use disorders in adult men and women with and without attention-deficit/hyperactivity disorder: A substantive and methodological overview. Neurosci. Biobehav. Rev. 151, 105209 (2023).

5. Krueger, R. F. & Eaton, N. R. Transdiagnostic factors of mental disorders. World Psychiatry 14, 27–29 (2015).

6. Linhartová, P., et al. Impulsivity in patients with borderline personality disorder: a comprehensive profile compared with healthy people and patients with ADHD. Psychol. Med. 50, 1829–1838 (2020).

7. Bénard, M., et al. Impulsivity is associated with food intake, snacking, and eating disorders in a general population. Am. J. Clin. Nutr. 109, 117–126 (2019).

8. de Wit, H. Impulsivity as a determinant and consequence of drug use: a review of underlying processes. Addict. Biol. 14, 22–31 (2009).

9. Van Dessel, J., et al. Waiting impulsivity: a distinctive feature of ADHD neuropsychology? Child Neuropsychol. 25, 122–129 (2019).

10. Marmorstein, N. R., White, H. R., Loeber, R. & Stouthamer-Loeber, M. Anxiety as a predictor of age at first use of substances and progression to substance use problems among boys. J. Abnorm. Child Psychol. 38, 211–224 (2010).

11. Wetherell, J. L., Gatz, M. & Pedersen, N. L. A longitudinal analysis of anxiety and depressive symptoms. Psychol. Aging 16, 187–195 (2001).

12. Kalin, N. H. The Critical Relationship Between Anxiety and Depression. Am. J. Psychiatry 177, 365–367 (2020).

13. Pedersen, W. Mental health, sensation seeking and drug use patterns: a longitudinal study. Br. J. Addict. 86, 195–204 (1991).

14. Hamdan-Mansour, A. M., Mahmoud, K. F., Al Shibi, A. N. & Arabiat, D. H. Impulsivity and Sensation-Seeking Personality Traits as Predictors of Substance Use Among University Students. J. Psychosoc. Nurs. Ment. Health Serv. 56, 57–63 (2018).

15. Nower, L., Derevensky, J. L. & Gupta, R. The relationship of impulsivity, sensation seeking, coping, and substance use in youth gamblers. Psychol. Addict. Behav. J. Soc. Psychol. Addict. Behav. 18, 49–55 (2004).

16. Ortin, A., Lake, A. M., Kleinman, M. & Gould, M. S. Sensation Seeking as Risk Factor for Suicidal Ideation and Suicide Attempts in Adolescence. J. Affect. Disord. 143, 214–222 (2012).

17. Daughters, S. B., et al. Distress Tolerance as a Predictor of Early Treatment Dropout in a Residential Substance Abuse Treatment Facility. J. Abnorm. Psychol. 114, 729–734 (2005).

18. Daughters, S. B., et al. Distress tolerance among substance users is associated with functional connectivity between prefrontal regions during a distress tolerance task. Addict. Biol. 22, 1378–1390 (2017).

19. Vujanovic, A. A., Bonn-Miller, M. O., Potter, C. M., Marshall, E. C. & Zvolensky, M. J. An Evaluation of the Relation Between Distress Tolerance and Posttraumatic Stress within a Trauma-Exposed Sample. J. Psychopathol. Behav. Assess. 33, 129–135 (2011).

20. Mattingley, S., Youssef, G. J., Manning, V., Graeme, L. & Hall, K. Distress tolerance across substance use, eating, and borderline personality disorders: A meta-analysis. J. Affect. Disord. 300, 492–504 (2022).

21. Watson, P., et al. Sign-tracking to non-drug reward is related to severity of alcohol-use problems in a sample of individuals seeking treatment. Addict. Behav. 154, 108010 (2024).

22. Anselme, P. & Robinson, M. J. F. From sign-tracking to attentional bias: Implications for gambling and substance use disorders. Prog. Neuropsychopharmacol. Biol. Psychiatry 99, 109861 (2020).

23. Versace, F., Kypriotakis, G., Basen-Engquist, K. & Schembre, S. M. Heterogeneity in brain reactivity to pleasant and food cues: evidence of sign-tracking in humans. Soc. Cogn. Affect. Neurosci. 11, 604–611 (2016).

24. Sha, Z., Wager, T. D., Mechelli, A. & He, Y. Common Dysfunction of Large-Scale Neurocognitive Networks Across Psychiatric Disorders. Biol. Psychiatry 85, 379–388 (2019).

25. Mohan, A., et al. The Significance of the Default Mode Network (DMN) in Neurological and Neuropsychiatric Disorders: A Review. Yale J. Biol. Med. 89, 49–57 (2016).

26. Goodkind, M., et al. Identification of a Common Neurobiological Substrate for Mental Illness. JAMA Psychiatry 72, 305–315 (2015).

27. Elsey, J. W. B., et al. Relationships between impulsivity, anxiety, and risk-taking and neural correlates of attention in adolescents. Dev. Neuropsychol. 41, 38–58 (2016).

28. Steele, J. S., et al. A specific neural substrate predicting current and future impulsivity in young adults. Mol. Psychiatry 26, 4919–4930 (2021).

29. Brown, M. R. G., et al. Neural correlates of high-risk behavior tendencies and impulsivity in an emotional Go/NoGo fMRI task. Front. Syst. Neurosci. 9, (2015).

30. Jung, H.-Y., Bak, H., Bang, M., Lee, S.-H. & Lee, K. S. Neural Correlates of Trait Impulsivity among Adult Healthy Individuals. Clin. Psychopharmacol. Neurosci. 22, 345–353 (2024).

31. Tozzi, L., et al. Personalized brain circuit scores identify clinically distinct biotypes in depression and anxiety. Nat. Med. 30, 2076–2087 (2024).

32. Xiao, X., et al. Brain Functional Connectome Defines a Transdiagnostic Dimension Shared by Cognitive Function and Psychopathology in Preadolescents. Biol. Psychiatry 95, 1081–1090 (2024).

33. Cruz, F. C., et al. Role of Nucleus Accumbens Shell Neuronal Ensembles in Context-Induced Reinstatement of Cocaine-Seeking. J. Neurosci. 34, 7437–7446 (2014).

34. Warren, B. L., et al. Separate vmPFC Ensembles Control Cocaine Self-Administration Versus Extinction in Rats. J. Neurosci. 39, 7394–7407 (2019).

35. Ramirez, S., et al. Activating positive memory engrams suppresses depression-like behaviour. Nature 522, 335–339 (2015).

36. Hammack, R. J., Fischer, V. E., Andrade, M. A. & Toney, G. M. Anterior basolateral amygdala neurons comprise a remote fear memory engram. Front. Neural Circuits 17, (2023).

37. Hammack, R. J., Fischer, V. E., Andrade, M. A. & Toney, G. M. Presence of a remote fear memory engram in the central amygdala. Learn. Mem. 30, 250–259 (2023).

38. Hope, B. T. Chapter 6 - Fos-Expressing Neuronal Ensembles in Addiction Research. in Neural Mechanisms of Addiction (ed. Torregrossa, M.) 75–88 (Academic Press, 2019). doi:10.1016/B978-0-12-812202-0.00006-3.

39. Liu, X., Wang, F., Le, Q. & Ma, L. Cellular and molecular basis of drug addiction: The role of neuronal ensembles in addiction. Curr. Opin. Neurobiol. 83, 102813 (2023).

40. Heilbronner, S. R., Rodriguez-Romaguera, J., Quirk, G. J., Groenewegen, H. J. & Haber, S. N. Circuit-Based Corticostriatal Homologies Between Rat and Primate. Biol. Psychiatry 80, 509–521 (2016).

41. Moschak, T. M. & Carelli, R. M. An opposing role for prelimbic cortical projections to the nucleus accumbens core in incubation of craving for cocaine versus water. Drug Alcohol Depend. 228, 109033 (2021).

42. West, E. A., et al. Noninvasive Brain Stimulation Rescues Cocaine-Induced Prefrontal Hypoactivity and Restores Flexible Behavior. Biol. Psychiatry 89, 1001–1011 (2021).

43. Ye, L., et al. Prelimbic cortex miR-34a contributes to (2*R*,6*R*)-hydroxynorketamine-mediated antidepressant-relevant actions. Neuropharmacology 208, 108984 (2022).

44. Khairuddin, S., et al. Prelimbic Cortical Stimulation Induces Antidepressant-like Responses through Dopaminergic-Dependent and -Independent Mechanisms. Cells 12, 1449 (2023).

45. Gao, F., et al. Elevated prelimbic cortex-to-basolateral amygdala circuit activity mediates comorbid anxiety-like behaviors associated with chronic pain. J. Clin. Invest. 133, (2023).

46. Jinks, A. L. & McGregor, I. S. Modulation of anxiety-related behaviours following lesions of the prelimbic or infralimbic cortex in the rat. Brain Res. 772, 181–190 (1997).

47. Sarica, C., et al. Prelimbic Cortex Deep Brain Stimulation Reduces Binge Size in a Chronic Binge Eating Rat Model. Stereotact. Funct. Neurosurg. 96, 33–39 (2018).

48. Blasio, A., Steardo, L., Sabino, V. & Cottone, P. Opioid system in the medial prefrontal cortex mediates binge-like eating. Addict. Biol. 19, 652–662 (2014).

49. Moschak, T. M. & Carelli, R. M. A sex-dependent role for the prelimbic cortex in impulsive action both before and following early cocaine abstinence. Neuropsychopharmacology 46, 1565–1573 (2021).

50. Narayanan, N. S., Cavanagh, J. F., Frank, M. J. & Laubach, M. Common medial frontal mechanisms of adaptive control in humans and rodents. Nat. Neurosci. 16, 1888–1895 (2013).

51. Smoak, M. A., Galvan, K. J., Calvo, D. E., Powers, R. E. & Moschak, T. M. Prelimbic Cortex Activity Predicts Anxiety-Like Behavior in the Elevated Plus Maze. Eur. J. Neurosci. 62, e70232 (2025).

52. Shimizu, T., Minami, C. & Mitani, A. Effect of electrical stimulation of the infralimbic and prelimbic cortices on anxiolytic-like behavior of rats during the elevated plus-maze test, with particular reference to multiunit recording of the behavior-associated neural activity. Behav. Brain Res. 353, 168–175 (2018).

53. Lanser, M. G., Ellenbroek, B. A., Zitman, F. G., Heeren, D. J. & Cools, A. R. The role of medial prefrontal cortical dopamine in spontaneous flexibility in the rat. Behav. Pharmacol. 12, 163 (2001).

54. Moschak, T. M., Sloand, T. J. & Carelli, R. M. Prelimbic Cortex Activity during a Distress Tolerance Task Predicts Cocaine-Seeking Behavior in Male, But Not Female Rats. J. Neurosci. Off. J. Soc. Neurosci. 43, 647–655 (2023).

55. Spring, M. G., Soni, K. R., Wheeler, D. S. & Wheeler, R. A. Prelimbic prefrontal cortical encoding of reward predictive cues. Synapse 75, e22202 (2021).

56. Spring, M. G., et al. Chronic Stress Prevents Cortico-Accumbens Cue Encoding and Alters Conditioned Approach. J. Neurosci. 41, 2428–2436 (2021).

57. Lovic, V., Saunders, B. T., Yager, L. M. & Robinson, T. E. Rats prone to attribute incentive salience to reward cues are also prone to impulsive action. Behav. Brain Res. 223, 255–261 (2011).

58. Chitre, A. S., et al. Genome-wide association study in a rat model of temperament identifies multiple loci for exploratory locomotion and anxiety-like traits. Front. Genet. 13, (2023).

59. Hughson, A. R. et al. Incentive salience attribution, “sensation-seeking” and “novelty-seeking” are independent traits in a large sample of male and female heterogeneous stock rats. Sci. Rep. 9, 2351 (2019).

60. Belin, D., Belin-Rauscent, A., Everitt, B. J. & Dalley, J. W. In search of predictive endophenotypes in addiction: insights from preclinical research. Genes Brain Behav. 15, 74–88 (2016).

61. Salamone, J. D. & Correa, M. Critical review of RDoC approaches to the study of motivation with animal models: effort valuation/willingness to work. Emerg. Top. Life Sci. 6, 515–528 (2022).

62. Moschak, T. M., Stang, K. A., Phillips, T. J. & Mitchell, S. H. Behavioral inhibition in mice bred for high vs. low levels of methamphetamine consumption or sensitization. Psychopharmacology (Berl.) 222, 353–365 (2012).

63. Moschak, T. M., Stang, K. A. & Mitchell, S. H. Mice Bred for Severity of Acute Alcohol Withdrawal Respond Differently in a Go/No-Go Task. Alcohol. Clin. Exp. Res. 37, 1483–1490 (2013).

64. Moschak, T. M., Terry, D. R., Daughters, S. B. & Carelli, R. M. Low distress tolerance predicts heightened drug seeking and taking after extended abstinence from cocaine self-administration. Addict. Biol. 23, 130–141 (2018).

65. Meyer, P. J., et al. Quantifying Individual Variation in the Propensity to Attribute Incentive Salience to Reward Cues. PLOS ONE 7, e38987 (2012).

66. Pnevmatikakis, E. A. et al. Simultaneous Denoising, Deconvolution, and Demixing of Calcium Imaging Data. Neuron 89, 285–299 (2016).

67. Giovannucci, A., et al. CaImAn an open source tool for scalable calcium imaging data analysis. eLife 8, e38173 (2019).

68. Moschak, T. M., Wang, X. & Carelli, R. M. A Neuronal Ensemble in the Rostral Agranular Insula Tracks Cocaine-Induced Devaluation of Natural Reward and Predicts Cocaine Seeking. J. Neurosci. 38, 8463–8472 (2018).

69. Wang, H.-T., et al. Finding the needle in a high-dimensional haystack: Canonical correlation analysis for neuroscientists. NeuroImage 216, 116745 (2020).

70. Hayes, A. F. Introduction to Mediation, Moderation, and Conditional Process Analysis, Second Edition: A Regression-Based Approach. (Guilford Publications, 2017).

71. Zuckerman, M. Behavioral Expressions and Biosocial Bases of Sensation Seeking. xiv, 463 (Cambridge University Press, New York, NY, US, 1994).

72. Bornovalova, M. A., et al. Risk taking differences on a behavioral task as a function of potential reward/loss magnitude and individual differences in impulsivity and sensation seeking. Pharmacol. Biochem. Behav. 93, 258–262 (2009).

73. Qianlan, Y., Shou, C., Tianya, H., Wei, D. & Liu, T. Sensation seeking and risk adjustment: the role of reward sensitivity in dynamic risky decisions. Front. Behav. Neurosci. 19, (2025).

74. Gray, J. A. A Critique of Eysenck’s Theory of Personality. in A Model for Personality (ed. Eysenck, H. J.) 246–276 (Springer, Berlin, Heidelberg, 1981). doi:10.1007/978-3-642-67783-0_8.

75. Wang, W. & Liu, H. Canonical correlation analysis of anxiety symptom and behavioral inhibition/activation system among college students and their relationship with physical activity. Sci. Rep. 15, 17547 (2025).

76. Grillon, C., et al. Clinical anxiety promotes excessive response inhibition. Psychol. Med. 47, 484–494 (2017).

77. Smillie, L. D., Jackson, C. J. & Dalgleish, L. I. Conceptual distinctions among Carver and White’s (1994) BAS scales: A reward-reactivity versus trait impulsivity perspective. Personal. Individ. Differ. 40, 1039–1050 (2006).

78. Espinoza Oyarce, D. A., Burns, R., Butterworth, P. & Cherbuin, N. Bridging Classical and Revised Reinforcement Sensitivity Theory Research: A Longitudinal Analysis of a Large Population Study. Front. Psychol. 12, (2021).

79. Heym, N., Ferguson, E. & Lawrence, C. An evaluation of the relationship between Gray’s revised RST and Eysenck’s PEN: Distinguishing BIS and FFFS in Carver and White’s BIS/BAS scales. Personal. Individ. Differ. 45, 709–715 (2008).

80. Crawford, J. L., Brough, R. E., Eisenstein, S. A., Peelle, J. E. & Braver, T. S. Generalized Encoding of the Relative Subjective Value of Cognitive Effort in the Dorsal ACC. J. Neurosci. 44, (2024).

81. Fernandez-Leon, J. A., et al. Neural correlates and determinants of approach–avoidance conflict in the prelimbic prefrontal cortex. eLife 10, e74950 (2021).

82. Capuzzo, G. & Floresco, S. B. Prelimbic and Infralimbic Prefrontal Regulation of Active and Inhibitory Avoidance and Reward-Seeking. J. Neurosci. 40, 4773–4787 (2020).

83. Iadipaolo, A. S., et al. Behavioral activation sensitivity and default mode network-subgenual cingulate cortex connectivity in youth. Behav. Brain Res. 333, 135–141 (2017).

84. Shinagawa, S., et al. Neural basis of motivational approach and withdrawal behaviors in neurodegenerative disease. Brain Behav. 5, e00350 (2015).

85. Zhang, Y., et al. Detailed mapping of behavior reveals the formation of prelimbic neural ensembles across operant learning. Neuron 110, 674–685.e6 (2022).

86. Gabriel, C. J., et al. Transformations in prefrontal ensemble activity underlying rapid threat avoidance learning. Curr. Biol. 35, 1128–1136.e4 (2025).

87. Cai, D. J., et al. A shared neural ensemble links distinct contextual memories encoded close in time. Nature 534, 115–118 (2016).

88. Giannotti, G., Heinsbroek, J. A., Yue, A. J., Deisseroth, K. & Peters, J. Prefrontal cortex neuronal ensembles encoding fear drive fear expression during long-term memory retrieval. Sci. Rep. 9, 10709 (2019).

89. Sortman, B. W., Gobin, C., Rakela, S., Cerci, B. & Warren, B. L. Prelimbic Ensembles Mediate Cocaine Seeking After Behavioral Acquisition and Once Rats Are Well-Trained. Front. Behav. Neurosci. 16, (2022).

90. Whitaker, L. R., et al. Bidirectional Modulation of Intrinsic Excitability in Rat Prelimbic Cortex Neuronal Ensembles and Non-Ensembles after Operant Learning. J. Neurosci. 37, 8845–8856 (2017).

91. Glanzberg, J. T., et al. Individual differences in prelimbic neural representation of food and cocaine seeking. Cell Rep. 43, (2024).

92. West, E. A., Saddoris, M. P., Kerfoot, E. C. & Carelli, R. M. Prelimbic and infralimbic cortical regions differentially encode cocaine-associated stimuli and cocaine-seeking before and following abstinence. Eur. J. Neurosci. 39, 1891–1902 (2014).

