## Supplement for "A shared ensemble in the prelimbic cortex links impulsivity and anxiety-like behavior"

**Supplemental Methods.**

Animal Housing and Surgery

Thirty-eight Sprague Dawley rats (19 males, 19 females) were obtained from Envigo (St. Charles, MO) at 8-10 weeks of age. Rats were individually-housed in ventilated polycarbonate cages with bedding in a temperature-controlled vivarium under a 12-hour reverse light/dark cycle (lights off at 08:00 AM). All behavioral testing occurred during the dark cycle. Rats had ad libitum access to water throughout the experiment. To facilitate motivation, food restriction was implemented during certain behavioral phases as noted below. All procedures were approved by the Institutional Animal Care and Use Committee (IACUC) of the University of Texas at El Paso and complied with NIH guidelines.

After arriving, rats were granted a 7 day acclimation period prior to surgery. Rats were induced with an anaesthetic cocktail (100mg/kg ketamine, 10mg/kg xylazine, IP) and maintained using 1–3% isoflurane (Covetrus) and/or supplemental anesthetic doses ("boosts") of the ketamine cocktail to maintain adequate anesthetic depth as assessed by pedal withdrawal. A midline scalp incision exposed the skull, and the skull surface was leveled. Four screws (two rostral, two caudal) were implanted into the skull, then a unilateral 2 mm circular craniotomy was drilled over the prelimbic cortex (AP: +2.7, ML: ±0.6) using a trephine burr. A 10 µL Hamilton syringe fitted with a 26-gauge beveled needle was then lowered to DV: –3.6 mm. A two minute rest was taken after the needle was lowered into position and before 700 nL of GCamp6S (AAV9-hSyn-GCaMP6s-WPRE-SV40, Addgene) was infused into the target site. After injection, the needle remained in place for 5 minutes.

Next, a 1.0mm diameter, 9.0 mm long GRIN lens (Inscopix) was implanted 200 µm above the injection site and secured with acrylic resin dental cement (Lang Dental). Animals received 5 mg/kg meloxicam (s.c.) and 10% enrofloxacin diluted in sterile saline and were monitored for post-operative recovery for at least 7 days before handling and behavioral testing resumed. 4–6 weeks following surgery the animals' GCAMP expression was verified utilizing the Miniscope software system (UCLA Miniscope). Once the presence of fluorescence of cells was observed, miniscope baseplates (OEPS, LABmaker) were affixed to the skull to support the miniscope camera attachment and in vivo recording during behavior tasks.

Apparatus

Five sound-attenuated behavioral chambers (16 in × 17 in × 27 in; Med Associates) were used for both the impulsivity and distress tolerance tasks. Each chamber was equipped with two retractable levers (Harvard Biosciences) positioned to the left and right of a centrally located pellet receptacle (Med Associates). A cue light was positioned above each lever, and levers were connected to a pellet dispensing mechanism (Med Associates). Pellets consisted of 45mg sucrose in either peanut-butter or raspberry flavor (TestDiet). A speaker was embedded above the cue light to emit an aversive white noise burst when incorrect responses were given.

For the Pavlovian Conditioned Approach (PAV) task, the chamber was structurally similar but differed in three ways: (1) the levers (Med Associates) were oriented vertically rather than horizontally, (2) the cue lights were embedded within the levers themselves instead of positioned above them, and (3) two pellet receptacles were present, each linked to a level and pellet dispenser, delivering the previously described peanut- or raspberry-flavored sucrose pellets.

Anxiety-like behavior was assessed using an Elevated Plus Maze (EPM; 44.5 in × 44.5 in × 44 in; Med Associates), consisting of a plus-shaped platform elevated from the floor, with two arms enclosed by high walls (“closed” arms) and two arms without enclosure (“open” arms) situated opposite each other. The EPM was connected to a Med PC control interface and desktop computer to allow for automated behavioral tracking.

Locomotor activity was measured in a square open field chamber (17 in × 17 in × 12 in; Med Associates) with a flat, smooth floor that allowed for unrestricted movement within a sound attenuating cubicle. Movement was tracked using infrared arrays that could measure X, Y, and Z positioning within the chamber.

All behavior responses, including lever presses, nose pokes, cue light illumination, and pellet delivery, were programmed and recorded using Med-PC IV software (Med Associates). Events were timestamped and synchronized with neural recordings.

Behavior

Following surgical recovery, rats underwent a battery of behavioral tasks designed to assess impulsivity, distress tolerance, Pavlovian conditioned approach (sign-tracking vs. goal-tracking), anxiety-like behavior, and locomotor activity. All behavioral sessions were recorded with the miniscope system, with behavior and calcium imaging synchronized using transistor-transistor logic (TTL). Animals were food-restricted to no less than 10 grams of Purina Laboratory Chow per day during the impulsivity and distress tolerance tasks and restricted to no less than 90% of their starting weight. Rats were subsequently put on ad libitum the day prior to the Elevated Plus Maze, Locomotor Activity, and Pavlovian Conditioning Approach tasks.

Impulsivity Task (IMP; **Fig. 1A**)

Rats first learned to press a lever for a sucrose pellet. After acquiring 50 reinforcers over two consecutive days, they then trained to respond within a 5 s window, cued by an illuminated light above the lever. Upon getting at least 50 rewards on the 5 s cue task, the animal began to train in the IMP task. The IMP task was composed of a variable (VT 1s) pre-cue period where the lever was extended, a cue period where the cue light was illuminated for a variable length of time (0.03 – 5 s depending on behavior, see below), and a fixed post-cue period (2s) where the lever stayed extended after the cue light had extinguished (as adapted from[1]). If the lever was pressed during the cue period (when the cue light was illuminated), a sucrose pellet was delivered. If the lever was pressed during either the pre-cue or post-cue periods, no pellet was delivered, and a brief white noise burst was emitted to indicate an “error.” Early errors were defined as lever presses occurring before cue light illumination (pre-cue period), and late errors as lever presses made after cue extinguishing (post-cue period). Impulsivity was quantified as the number of early errors divided by the total number of lever presses. The duration of the cue period varied based on each animal’s previous performance (correct responses decreased the cue period, incorrect responses lengthened the cue period). At the start of each session, the cue duration was initialized to the final cue duration from the preceding session. Each session continued until either 100 lever presses were recorded or 1 hour had passed.

Distress Tolerance Task (DT; **Fig. 1B**)

The distress tolerance task was adapted from[2]. The DT task was recorded following the IMP task and the structure was similar with a pre-cue period, cue period, and post-cue period.

However, unlike the IMP task, the cue light duration progressively decreased following each response until it was impossible for the rat to obtain a correct response. The initial cue duration was individualized for each animal based on prior task performance (see[2]). Lever presses during the cue period resulted in pellet delivery, while responses outside the cue period triggered a brief white noise burst. Distress tolerance was defined as the duration in the task before the rat gave up ('giving up' was operationally defined as having 8 out of 10 responses that are omissions with at least 5 consecutive omissions).

Pavlovian Conditioned Approach (PAV; **Fig. 1C**)

Rats underwent a 5-day PCA training protocol prior to recording. Each session consisted of 25 trials with a 90 s variable inter-trial interval (ITI). On each trial, the retractable lever was extended for 8 seconds, followed immediately by pellet delivery into the corresponding receptacle. No response was required from the rat to receive the pellet.

During the lever presentation, lever interactions and head entries into the food cup were recorded. A Pavlovian conditioned approach index was calculated to quantify sign-tracking or goal-tracking preference using approach probability, response bias, and initial preference (modified from[3]). Approach probability was defined as the difference between the probability of interacting with the lever and the probability of interacting with the food cup, response bias was defined as the ratio of lever interactions and food cup interactions over the total number of interactions, and initial preference was defined as the proportion of trials with a lever interaction first minus the proportion of trials with a food cup interaction first. Averaging these three measures yielded a Pavlovian conditioned approach score ranging from -1 (goal-tracking) to 1 (sign-tracking). Pellet flavor in the PCA task was counterbalanced to differ from the flavor used in the impulsivity and distress tolerance tasks.

Anxiety-like behavior (ANX; **Fig. 1D**)

The elevated plus maze was used to assess anxiety-like behavior. The test room was a dimly lit, open area with the maze centrally located. Once the test was initiated, the researcher would leave the animal alone in the silent room. Animals were placed in the direct center of the maze, facing an open arm and allowed to explore for 10 minutes. Behavior was recorded via the number of infrared beam breaks at the junction of each arm. Metrics analyzed included percent time spent in the open arms, number of entries into each arm, and total distance traveled. Greater open-arm time was interpreted as reduced anxiety-like behavior. After the initial 10 minutes, the animal was removed from the maze and returned to their home cage for another 10 minutes to allow a larger duration of recording time to better isolate neural units. Importantly, a subset of the elevated plus maze data we describe here was previously documented in a short report[4]; this is specifically relevant to **Fig. 2L**. However, we significantly expand upon these findings in the remaining figures and **Fig. 2L** itself is neither identical to figures in our prior report nor central to the premise of the current manuscript. Two animals spent less than 20s in the open arms and were thus excluded from the neural analysis as there was too little activity to accurately analyze.

Locomotor Activity (LOCO; **Fig. 1E**)

Locomotor activity was measured in a square open-field arena with opaque walls under dim light. Animals were allowed to explore the arena freely for 10 minutes. Activity Monitor software

(Med Associates) monitors movements and positioning within the arena. Total distance travelled was recorded as our measure of interest.

Histology

At study completion, rats were deeply anesthetized with 5% isoflurane and euthanized by decapitation. Brains were extracted and fixed in 4% paraformaldehyde for 24 hours, followed by cryoprotection in 30% sucrose for at least 3 days. Coronal sections (30 μm) were cut using a Leica CM3050 S cryostat. Sections were wet-mounted onto microscope slides and imaged using a Zeiss LSM 900 confocal microscope to verify GRIN lens placement utilizing the Paxinos and Watson (2007) brain atlas (See **Fig. S4**).

Calcium Imaging (**Fig. 2A-B**)

In vivo calcium imaging was recorded during behavioral tasks using the UCLA Miniscope endoscopic camera system. During each task, the camera was attached to the implanted baseplate on the animal's skull and removed immediately after the task was over. Behavioral events programmed in Med-PC (v.5) were synchronized to the calcium imaging data via a transistor-transistor logic (TTL) connection between Med-PC and the Miniscope software on a Dell laptop. Recordings were captured continuously throughout each behavioral task, with 5-minute breaks every 10 minutes to reduce the risk of photobleaching. These breaks did not affect behavioral timing.

Miniscope recordings were manually reviewed for quality and checked for any motion disturbances caused by camera shifts from head or body movements. Raw video files were downsampled, merged by timestamp if from the same session, and processed using CaImAn (Pnevmatikakis et al., 2016; Giovannucci et al., 2019). Motion correction was performed using CaImAn's built-in NoRMCorre algorithm. Putative neurons were identified using a deconvolution algorithm that highlights candidate neurons based on the spatial/temporal signature of each calcium signal. All candidate neurons were then manually inspected, and any that appeared to be artifacts or noise were excluded. Neurons were subsequently coregistered across each pair of tasks using CaImAn's RegisterMulti algorithm. Neural data were then aligned with behavioral event timestamps using MATLAB and Neuroexplorer for subsequent analyses.

Data analysis.

*Neural Activity and Classification*. Neurons were classified according to their activity in response to task events, similar to what we have done previously[1,4,5]. Baseline for each peri-event histogram was defined as the average activity in the 10 s prior to the event and data were normalized (z-score) to this baseline. Bonferroni-corrected paired t-tests compared baseline to the three 200 ms bins (600 ms total) following the relevant event. Neurons were classified as either "Excited" (significant increase in activity), "Inhibited" (significant decrease in activity), or "Nonphasic" (no change in activity) with respect to the event in question. For the impulsivity, distress tolerance, and Pavlovian conditioned approach tasks our primary focus was on the extension of the lever into the chamber (signaling trial initiation), based on ours and others' previous work with these tasks[1,6,7]. For each of these tasks, we then categorized "Excited" or "Inhibited" neurons according to the animals' subsequent behavior following trial initiation. For the impulsivity and distress tolerance tasks, animals could make an early error, a correct response, a late error or an omission. For the Pavlovian conditioned approach task, animals could make a lever response, a nosepoke response, or no response. For impulsivity, we examined the change in neural activity (specifically the average of the 600 ms after lever extension) between trials where the animal made an early error minus trials where the animal made a late error, as this tracked impulsive vs. non-impulsive responding. For distress tolerance, we examined the change in neural activity between trials where the animal made an error response (either early or late) minus trials where the animal made an omission, as this tracked high vs. low persistence in the task (errors were used rather than correct responses or total responses because there were very few correct responses and they came early on in the task). For Pavlovian conditioned approach, we examined the change in neural activity between trials where the animal made a lever response minus trials where the animal made a nosepoke response, as this tracked sign-tracking vs. goal-tracking. We then took a weighted average of these changes in neural activity for excited and inhibited neurons for each rat and each behavior, where inhibited activity was multiplied by -1 to account for its opposing direction. Thus, each rat had a single neural data point for each behavior. We used a different type of analysis for elevated plus maze and locomotor activity, as they did not have a trial-based structure. For elevated plus maze, we calculated the relative neural activity per zone (open arms, closed arms, junction) over the total activity for the session as a measure of "Neural preference" (e.g., open arm activity / total activity) as we did previously[4]. For locomotor activity we calculated the relative neural activity when animals had "locomotion" vs. when animals had "no locomotion". "Locomotion" was classified as each 1s bin where the animal moved at least 5 cm; "No locomotion" was all other bins. For both elevated plus maze and locomotor activity, we then took the average activity for each rat to give a single data point for each behavior. Thus, for all subsequent analysis, each animal had 1 behavioral and 1 neural data point for each behavior. All statistical outliers were winsorized[8] and all data were normalized before entering them into subsequent analyses.

2.1% of behavioral data and 15.2% of neural data were missing across all tasks. Thus, for subsequent principal component analyses (PCA) and canonical correlation analyses (CCA) these data were estimated. For behavioral data, we ran a linear regression between the behavioral data points in this study and a second behavioral data set from the same animals that were part of a greater longitudinal study beyond the scope of this manuscript. Since all missing data points for our study had existing data points for the second data set, we were able to use the slope of the linear regression to estimate the missing behavioral data points. For the neural data, we ran a linear regression between behavior and neural activity. Using this slope of this line, we then estimated the missing neural activity data points. Importantly, after completing PCA and CCA we reported the correlations between the raw data points *without* estimated data to the principal components and canonical variates we obtained to ensure the estimated data points were not artificially biasing the data.

PCA with varimax rotation was conducted on all behavioral and neural data (3 separate analyses for males, females, and combined; **Figs. 1E *top*, 2A *top*, 3C *top***). Concomitant with the reported loadings (which necessarily include estimated data), we also report the correlations of each principal component with each behavior *without* estimated data (**Figs. 1E *bottom*, 2A *bottom*, 3C *bottom***).

To investigate the relationship between shared neural and shared behavioral activity, we used CCA as per[9] (**Figs. 3B, 4D, S2A**). Individual regression scores from extracted neural principal components were input as X, while individual regression scores from extracted behavioral principal components were input as Y. We used a permutation test (10,000 permutations) followed by a Benjamini-Hochberg correction for multiple comparisons to assess significance for each pair of canonical variates. Subsequently we estimated the contribution of each principal component to the canonical variate (**Figs. 3C, S2B, S3**), and used Pearson correlations to compare the canonical variates with raw neural activity and raw behavioral data for each behavior, excluding estimated values (**Figs. 3D, 4F, S2C**). Importantly, for PCA and CCA analysis, we sometimes switched (**Fig. 3A *middle***) or inverted (**Fig. 4F**) axes when presenting the data to allow for easier comparison of findings across groups. In the case of **Fig. 3A *middle*** (male rats neural PCA), we felt this was justified because axes are determined solely by which principal component explains the most variance, but we wanted to stress the similarity in structure across groups. PC1 in the males very clearly aligns with PC2 in the overall analysis and in females, so we felt justified in making the switch for visual comparison. In the case of **Fig. 4F**, the direction (positive or negative) of the variates of the CCA is arbitrary. Thus, to more easily visualize the comparison between **Fig. 3D** and **4F**, we felt the inversion of the y-axis was justified.

We were also interested in neurons that either did or did not share patterns of neural activity across behaviors, focusing on those neurons that coregistered across 2 or more behaviors. To do so, we focused on the four behaviors wherein PL activity predicted behavior, thus excluding locomotor activity. To determine if common patterns existed across neurons, we first classified all neurons as “Excited”, “Inhibited” or “Nonphasic” for the IMP, DT, and PAV tasks, as described above. For each pair of behaviors in each rat, we calculated the percent of phasic neurons that had the same sign (e.g. if a neuron was “Inhibited” in both behaviors) minus the percent of phasic neurons that had opposing signs (e.g. if a neuron was “Inhibited” in one task but “excited” in another). For comparisons with ANX (which did not have a trial-based structure that could allow a similar excited/inhibited classification), we first established relative open arm activity for each neuron. Then, each neuron was classified as “excited” or “inhibited” based on activity in the other task to be compared (i.e. if we were comparing IMP and ANX, each ANX neuron would be sorted by whether it was excited or inhibited in the IMP task). Finally, we subtracted the relative open arm activity from neurons that were inhibited in the comparator task from the relative open arm activity from neurons that were excited in the comparator task. Data from all task comparisons were then assessed using a 1-sample t-test vs. 0 (0 being the point at which there is no difference between the percent of same vs opposing sign neurons). Thus, each pair of behaviors could reveal a positive dominant pattern of “neural preference” (same sign), a negative dominant pattern of “neural preference” (opposing signs) or no dominant pattern (**Fig. 4A,B**). For all significant t-tests, neurons that followed the dominant pattern were considered “shared”, while those that did not were considered “unshared”. To be able to classify ANX neurons, neurons with relative activity in the top quartile were considered “Excited”, bottom quartile “Inhibited” and all remaining “Nonphasic”. This allowed us to determine if individual ANX neurons could be considered “shared” or “unshared” with other tasks. We then calculated neural activity for each behavior and rat separately for shared and unshared neurons. For behaviors that shared activity with more than one other behavior, neural activity was averaged per rat. We thereafter ran PCA and CCA analysis on shared and unshared neurons, respectively (**Fig.4C,D**). Shared and unshared PCA and CCA analyses both used the same locomotor neural activity values from the original analysis, as these were not split into shared and unshared groups (we retained locomotor neural activity to allow comparison to the overall PCA and CCA analyses). Finally, we compared shared and unshared

CCA analysis to the overall CCA analysis by correlating the respective U and V scores and comparing the correlations using a Fisher z-transformation (**Fig. 4E**).

We next assessed if individual differences in the proportion of shared neurons between two given tasks predicted the proportion of shared neurons between two other tasks using Pearson correlations (**Fig. 4G**). Subsequently, we calculated a “neural overlap” score across each rat by averaging their “neural preference” (defined above) across all behaviors that either showed a statistically significant dominant pattern or that significantly correlated with another pattern (here, all lever based tasks as well as the pairing between impulsivity and elevated plus maze). Next, we used the moderation analysis macro PROCESS[10] to use an OLS regression model to investigate how “neural overlap” moderated the relationship between IMP and behavioral PC2 as well as between IMP and ANX.

**Supplemental Tables.**

**Table S1. Sex Differences.**

| **Task** | ***t*** | ***df*** | **p** |
|---|---|---|---|
| Impulsivity | 0.6417 | 36 | 0.5251 |
| Distress Tolerance | 1.4087 | 36 | 0.1675 |
| Pavlovian Conditioned Approach | 0.5791 | 33 | 0.5665 |
| Anxiety-like behavior | 0.3708 | 36 | 0.7129 |
| Locomotor Activity | 1.4128 | 35 | 0.1666 |

**Table S2. Principal components correlation with raw behavioral data.**

| **All rats.** | **PC1** | | **PC2** | |
|---|---|---|---|---|
| **Task** | ***r*** | **p** | ***r*** | **p** |
| Impulsivity | -0.127 | 0.449 | 0.887 | <0.001 |
| Distress Tolerance | 0.654 | <0.001 | 0.102 | 0.541 |
| Pavlovian Conditioned Approach | 0.525 | 0.001 | -0.063 | 0.721 |
| Anxiety-like behavior | -0.393 | 0.015 | -0.676 | <0.001 |
| Locomotor Activity | 0.792 | <0.001 | 0.174 | 0.304 |

| **Males.** | **PC1** | | **PC2** | |
|---|---|---|---|---|
| **Task** | ***r*** | **p** | ***r*** | **p** |
| Impulsivity | -0.006 | 0.980 | 0.874 | <0.001 |
| Distress Tolerance | 0.866 | <0.001 | 0.044 | 0.859 |
| Pavlovian Conditioned Approach | 0.742 | <0.001 | 0.093 | 0.723 |
| Anxiety-like behavior | -0.343 | 0.150 | -0.803 | <0.001 |
| Locomotor Activity | 0.683 | 0.001 | 0.431 | 0.065 |

| **Females.** | **PC1** | | **PC2** | |
|---|---|---|---|---|
| **Task** | ***r*** | **p** | ***r*** | **p** |
| Impulsivity | -.449 | 0.054 | 0.578 | 0.009 |

| | | | | |
|---|---|---|---|---|
| Distress Tolerance | 0.519 | 0.023 | 0.597 | 0.007 |
| Pavlovian Conditioned Approach | 0.039 | 0.878 | -0.710 | <0.001 |
| Anxiety-like behavior | 0.366 | 0.123 | 0.009 | 0.970 |
| Locomotor Activity | 0.815 | <0.001 | -0.073 | 0.773 |

**Table S3. Principal components correlation with raw neural data.**

| **All rats.** | **PC1** | | **PC2** | |
|---|---|---|---|---|
| **Task** | ***r*** | **p** | ***r*** | **p** |
| Impulsivity | 0.739 | <0.001 | -0.150 | 0.420 |
| Distress Tolerance | 0.855 | <0.001 | 0.119 | 0.524 |
| Pavlovian Conditioned Approach | 0.067 | 0.757 | -0.732 | <0.001 |
| Anxiety-like behavior | -0.109 | 0.628 | 0.774 | <0.001 |
| Locomotor Activity | 0.428 | 0.014 | 0.676 | <0.001 |

| **Males.** | **PC1** | | **PC2** | |
|---|---|---|---|---|
| **Task** | ***r*** | **p** | ***r*** | **p** |
| Impulsivity | 0.026 | 0.917 | 0.927 | <0.001 |
| Distress Tolerance | -0.148 | 0.584 | 0.158 | 0.560 |
| Pavlovian Conditioned Approach | -0.787 | 0.002 | 0.131 | 0.685 |
| Anxiety-like behavior | 0.717 | 0.013 | -0.491 | 0.125 |
| Locomotor Activity | 0.757 | <0.001 | 0.229 | 0.360 |

| **Females.** | **PC1** | | **PC2** | |
|---|---|---|---|---|
| **Task** | ***r*** | **p** | ***r*** | **p** |
| Impulsivity | 0.805 | <0.001 | 0.021 | 0.946 |
| Distress Tolerance | 0.892 | <0.001 | 0.231 | 0.408 |
| Pavlovian Conditioned Approach | 0.103 | 0.751 | -0.818 | 0.001 |
| Anxiety-like behavior | -0.039 | 0.908 | 0.449 | 0.165 |
| Locomotor Activity | 0.471 | 0.089 | 0.772 | 0.001 |

**Table S4. Principal components correlation with raw neural data.**

| **Shared Neurons.** | **PC1** | | **PC2** | |
|---|---|---|---|---|
| **Task** | ***r*** | **p** | ***r*** | **p** |
| Impulsivity | 0.768 | <0.001 | -0.211 | 0.310 |
| Distress Tolerance | 0.822 | <0.001 | 0.216 | 0.279 |
| Pavlovian Conditioned Approach | 0.665 | 0.007 | -0.090 | 0.750 |
| Anxiety-like behavior | -0.421 | 0.197 | 0.784 | 0.004 |
| Locomotor Activity | 0.216 | 0.290 | 0.884 | <0.001 |

| **Unshared Neurons.** | **PC1** | | **PC2** | |
|---|---|---|---|---|
| **Task** | ***r*** | **p** | ***r*** | **p** |
| Impulsivity | -0.110 | 0.586 | 0.197 | 0.325 |

| | | | | |
|---|---|---|---|---|
| Distress Tolerance | 0.915 | <0.001 | 0.154 | 0.461 |
| Pavlovian Conditioned Approach | -0.148 | 0.646 | 0.892 | <0.001 |
| Anxiety-like behavior | 0.215 | 0.424 | -0.761 | <0.001 |
| Locomotor Activity | 0.827 | <0.001 | -0.353 | 0.077 |

**Table S5. Canonical correlates correlation with raw data.**

| **Shared Neurons.** | **Neural** | | **Behavioral** | |
|---|---|---|---|---|
| **Task** | ***r*** | **p** | ***r*** | **p** |
| Impulsivity | -0.449 | 0.019 | -0.883 | <0.001 |
| Distress Tolerance | -0.303 | 0.126 | -0.151 | 0.455 |
| Pavlovian Conditioned Approach | -0.501 | 0.029 | -0.098 | 0.649 |
| Anxiety-like behavior | 0.525 | 0.037 | 0.866 | <0.001 |
| Locomotor Activity | 0.433 | 0.027 | -0.302 | 0.134 |

| **Unshared Neurons.** | **Neural** | | **Behavioral** | |
|---|---|---|---|---|
| **Task** | ***r*** | **p** | ***r*** | **p** |
| Impulsivity | -0.378 | 0.052 | -0.061 | 0.766 |
| Distress Tolerance | -0.433 | 0.024 | -0.717 | <0.001 |
| Pavlovian Conditioned Approach | -0.596 | 0.007 | -0.587 | 0.003 |
| Anxiety-like behavior | 0.072 | 0.788 | 0.500 | 0.008 |
| Locomotor Activity | 0.024 | 0.909 | -0.833 | <0.001 |

**Supplemental Figures.**

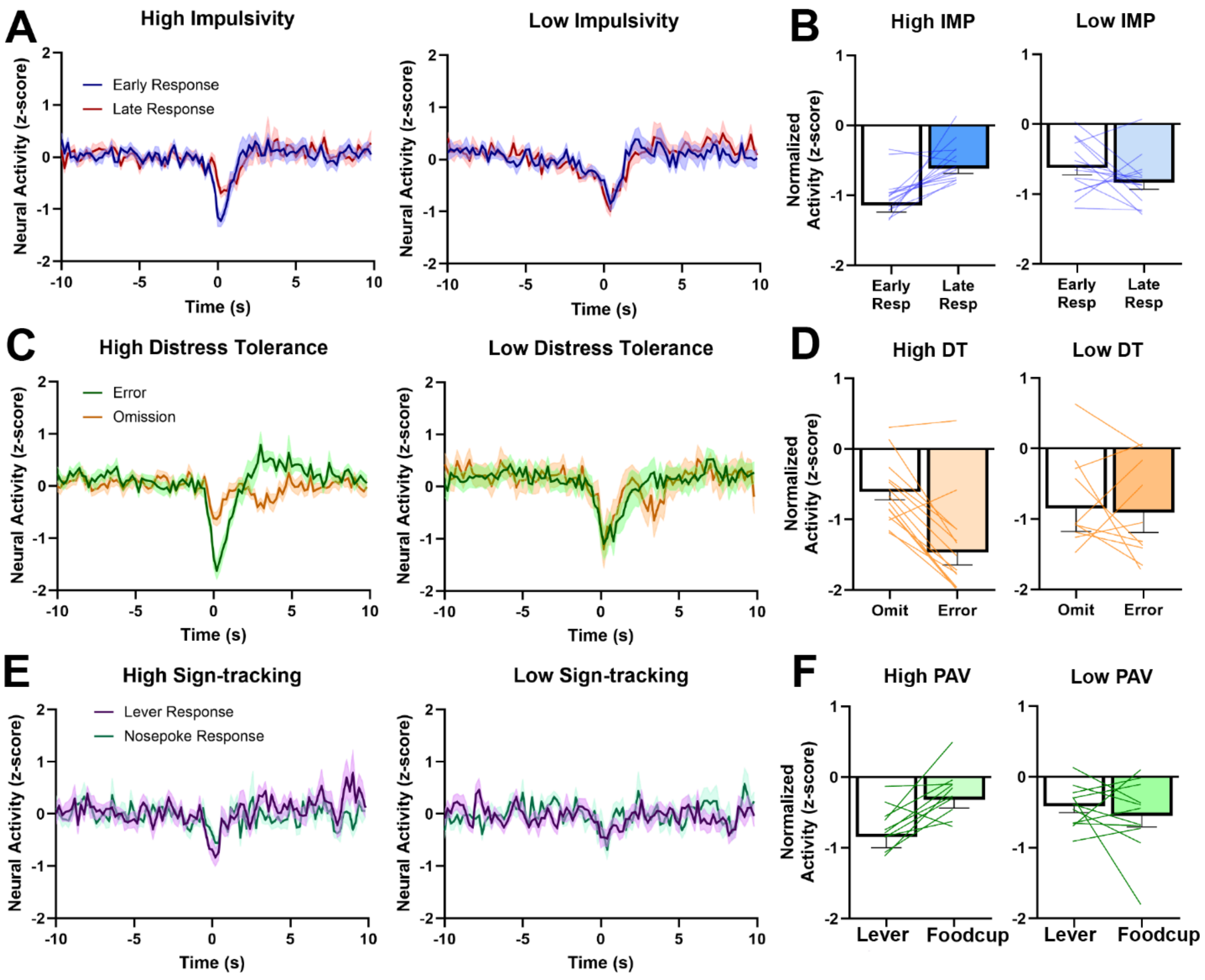


**Supplemental Figure 1.** Inhibited prelimbic activity during behavior. **A)** PL activity in neurons that were inhibited by trial initiation in the impulsivity task. Animals with low impulsivity had a more pronounced decrease in PL activity when they subsequently chose a late response, while animals with high impulsivity showed the opposite pattern. This pattern is statistically depicted in (**B**). **C)** PL activity in neurons that were inhibited by trial initiation in the distress tolerance task. Animals with high distress tolerance had a more pronounced decreases in PL activity when they subsequently made a response, while animals with low distress tolerance did not. This pattern is statistically depicted in (**D**). **E)** PL activity in neurons that were inhibited by trial initiation in the Pavlovian conditioned approach task. Animals with high sign-tracking had a more pronounced decrease in PL activity when they subsequently approached the lever, while animals with low sign-tracking showed the opposite pattern. This pattern is statistically depicted in (**E**). * p < 0.05, ** p < 0.01, *** p < 0.001.

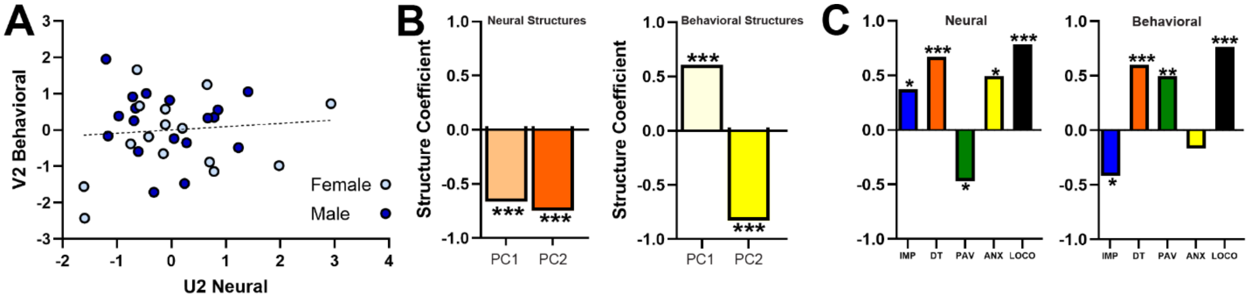


**Supplemental Figure 2.** Our canonical correlation analysis yielded two pairs of variates, $U_1$ and $V_1$ (presented in **Fig. 3B-D**), and $U_2$ and $V_2$, which we present in this figure. **A)** $U_2$ and $V_2$ did not significantly correlate following CCA. **B)** The two neural PCs both played a role in the shared variance of $U_2$, as each had a significant negative impact on the shared neural activity (PC1: $r = -0.663$, $p < 0.001$; PC2: $r = -0.7483$, $p < 0.001$; *left*). Conversely, the behavioral PCs had significant opposing impacts on the shared behavioral activity in $V_2$ (PC1: $r = 0.607$, $p < 0.001$; PC2: $r = -0.83$, $p < 0.001$; *right*). **C)** Our neural canonical variate ($U_2$) positively correlated with neural activity in the IMP, DT, and PAV tasks while correlating negatively with neural activity in the ANX task (IMP: $r = 0.373$, $p = 0.039$; DT: $r = 0.671$; $p < 0.001$; PAV: $r =$ **-** $0.472$, $p = 0.020$; ANX: $r = 0.493$, $p = 0.020$; LOCO: $r = 0.784$, $p < 0.001$; **Fig. 3D *left***). Our behavioral canonical variate ($V_1$) had a strong positive association with IMP and a strong negative association with ANX, as well as a weaker positive association with LOCO (IMP: $r = 0.412$, $p = 0.015$; DT: $r = 0.600$; $p < 0.001$; PAV: $r = 0.496$, $p = 0.005$; ANX: $r = -0.168$, $p = 0.350$; LOCO: $r = 0.762$, $p < 0.001$; **Fig. 3D *right***).

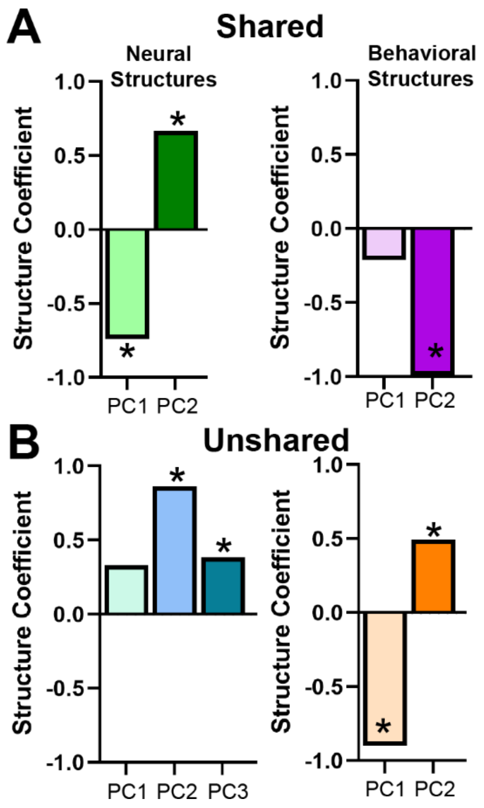


**Supplemental Figure 3.** In addition to our comparisons of the shared and unshared CCA to the overall CCA (**Fig. 4E**), we also determined how the individual principal components contributed to the structure of the shared and unshared CCA (similar to that done in **Fig. 3C**). The shared neuron analysis exhibited a very similar structure to the overall analysis: Neural PC1 and PC2 both significantly and oppositely contributed to the neural variate $U_1$ (**A *left***; Neural $PC1_{shared}$: r = -0.743 , p < 0.001; Neural $PC2_{shared}$: r = 0.669, p < 0.001), while only behavioral PC2 contributed to the behavioral variate $V_1$ (**A *right***; Behavioral $PC1_{shared}$: r = -0.221 , p = 0.291; Behavioral $PC2_{shared}$: r = -0.990 , p < 0.001; note that the contribution for each structure is inverted compared to the overall analysis in **Fig. 3C**. However, the direction of the signs [positive or negative] is arbitrary in CCA and does not indicate an inverted relationship). Conversely, the structure of the unshared neuron analysis was very different from the structure of the overall analysis. Notably, there were three neural PCs rather than two, and while two of them contributed to the structure of neural variate $U_1$ they did not contribute in an opposing manner (**B *left***; Neural $PC1_{unshared}$: r = 0.332, p = 0.090; Neural $PC2_{unshared}$: r = 0.861, p < 0.001; Neural $PC3_{unshared}$: r = 0.385, p = 0.047). Further, unlike the overall analysis, both behavioral PCs contributed to the structure of behavioral variate $V_1$ (**B *right***; Behavioral $PC1_{unshared}$: r = 0.900, p < 0.001; Behavioral $PC2_{unshared}$: r = -0.496, p = 0.009).

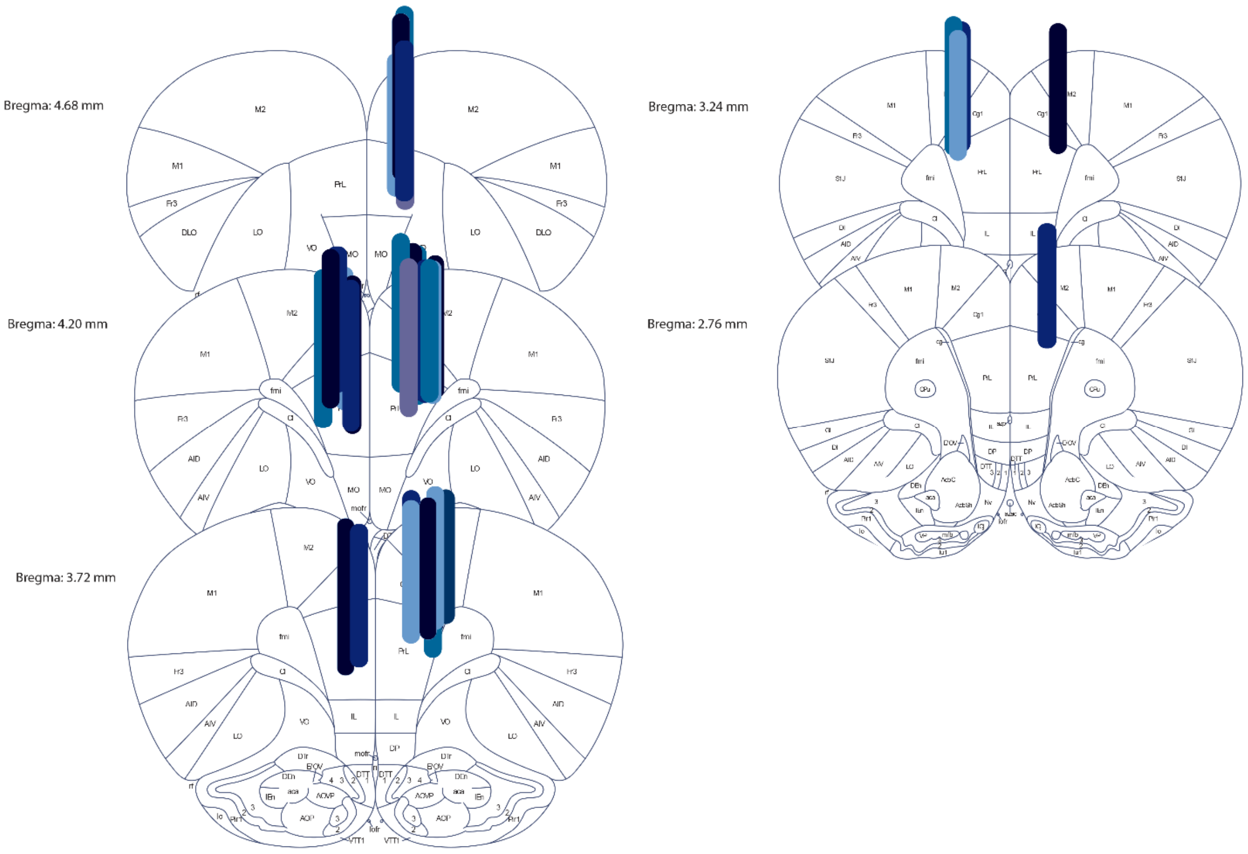


**Supplemental Figure 4.** Coronal maps displaying the placements of lenses in the prelimbic cortex.

**Supplemental References.**


1. Moschak, T. M. & Carelli, R. M. A sex-dependent role for the prelimbic cortex in impulsive action both before and following early cocaine abstinence. *Neuropsychopharmacol.* **46**, 1565–1573 (2021).

2. Moschak, T. M., Terry, D. R., Daughters, S. B. & Carelli, R. M. Low distress tolerance predicts heightened drug seeking and taking after extended abstinence from cocaine self-administration. *Addiction Biology* **23**, 130–141 (2018).

3. Meyer, P. J. *et al.* Quantifying Individual Variation in the Propensity to Attribute Incentive Salience to Reward Cues. *PLOS ONE* **7**, e38987 (2012).

4. Smoak, M. A., Galvan, K. J., Calvo, D. E., Powers, R. E. & Moschak, T. M. Prelimbic Cortex Activity Predicts Anxiety-Like Behavior in the Elevated Plus Maze. *European Journal of Neuroscience* **62**, e70232 (2025).

5. Moschak, T. M., Wang, X. & Carelli, R. M. A Neuronal Ensemble in the Rostral Agranular Insula Tracks Cocaine-Induced Devaluation of Natural Reward and Predicts Cocaine Seeking. *J Neurosci* **38**, 8463–8472 (2018).

6. Moschak, T. M., Sloand, T. J. & Carelli, R. M. Prelimbic Cortex Activity during a Distress Tolerance Task Predicts Cocaine-Seeking Behavior in Male, But Not Female Rats. *J Neurosci* **43**, 647–655 (2023).

7. Spring, M. G. *et al.* Chronic Stress Prevents Cortico-Accumbens Cue Encoding and Alters Conditioned Approach. *J. Neurosci.* **41**, 2428–2436 (2021).

8. Ruppert, D. Trimming and Winsorization. in (2014). doi:10.1002/9781118445112.stat01887.

9. Wang, H.-T. *et al.* Finding the needle in a high-dimensional haystack: Canonical correlation analysis for neuroscientists. *NeuroImage* **216**, 116745 (2020).

10. Hayes, A. F. *Introduction to Mediation, Moderation, and Conditional Process Analysis, Second Edition: A Regression-Based Approach*. (Guilford Publications, 2017).
